# Glycans function as a Golgi export signal to promote the constitutive exocytic trafficking

**DOI:** 10.1101/2020.05.20.105544

**Authors:** Xiuping Sun, Hieng Chiong Tie, Bing Chen, Lei Lu

**Affiliations:** School of Biological Sciences, Nanyang Technological University, 60 Nanyang Drive, Singapore 637551

**Keywords:** Golgi, Golgi export, O-glycosylation, N-glycosylation, secretory pathway, protein trafficking (Golgi), membrane trafficking, intracellular trafficking, membrane protein

## Abstract

Most proteins in the secretory pathway are glycosylated. However, the role of glycans in the membrane trafficking is still unclear. Here, we discovered that transmembrane secretory cargos, such as interleukin 2 receptor α subunit or Tac, transferrin receptor and cluster of differentiation 8a, unexpectedly displayed substantial Golgi localization when their O-glycosylation was compromised. By quantitatively measuring their Golgi residence times, we found that the apparent Golgi localization of these O-glycan deficient cargos is due to their slow Golgi export. The super-resolution microscopy method that we previously developed revealed that O-glycan deficient Tac chimeras localize at the interior of the *trans*-Golgi cisternae. The O-glycan was observed to be both necessary and sufficient for the efficient Golgi export of Tac chimeras. By sequentially introducing O-glycosylation sites to β-galactoside α-2,6-sialyltransferase1, we demonstrated that the O-glycan’s effect on the Golgi export is probably additive. Finally, the finding that N-glycosylated GFP substantially reduces the Golgi residence time of Tac chimera suggests that the N-glycan might have a similar effect. Therefore, both O- and N-glycan might function as a generic Golgi export signal at the *trans*-Golgi to promote the constitutive exocytic trafficking.

## Introduction

In the secretory pathway, newly synthesized membrane proteins (cargos) are exported from the endoplasmic reticulum (ER) toward the Golgi, where they sequentially pass through cisternae of the Golgi stack. Once reaching the *trans*-side of the Golgi, cargos are packed into membrane carries destined for the plasma membrane (PM) or endolysosomes (1-3). It is well accepted that protein trafficking is mediated by a signal, usually a cytosolic short stretch of amino acids (AAs) that is recognized by diverse trafficking machineries. In the secretory pathway, the efficient ER export of many cargos can be facilitated by COPII-coat-binding ER export signals (4,5). At the *trans*-side of the Golgi, secretory cargos that possess clathrin adaptor interacting signals are targeted to the endolysosome in clathrin-coated vesicles (1-3,6). Without such targeting or export signals, the majority of secretory cargos, such as vesicular stomatitis glycoprotein G (VSVG), follow the by default constitutive exocytic pathway and are exported from the Golgi in non-coated membrane carriers (7-9). So far, the generic Golgi export signal has not been identified for the constitutive pathway, though a putative signal was reported in a viral membrane protein (10). Lacking a conventional coat-like machinery, it is also unclear how these cargos are sorted from Golgi transmembrane resident proteins (hereafter residents) and recruited to the exocytic membrane carriers.

Almost all secretory proteins are glycosylated in the ER and/or Golgi. However, the roles of polysaccharide chains or glycans in the protein secretion are still unclear. The best-known example is the mannose 6-phosphate (M6P)-modified glycan which specifically attaches to the soluble hydrolytic enzymes of the lysosome (11,12). At the TGN, M6P is recognized by M6P receptor, which subsequently concentrates lysosomal enzymes to clathrin-coated vesicles targeting to the endolysosome. As another example, at the ER exit site, the cargo receptor, ERGIC53, binds to the high-mannose N-glycans of several soluble secretory proteins, such as coagulation factor V and VII, for their efficient ER export through COPII-coated vesicles (13,14).

Although the molecular mechanism behind is still obscure, available data indicate that glycans are also involved in the sorting and trafficking of secretory cargos to the PM. In cargos without basolateral PM sorting signals, glycans have been demonstrated by multiple research groups to act as an apical PM targeting signal in polarized epithelial cells (15,16). Notably, an attempt has been made by Gut *et al*. on the role of the N-glycosylation in the Golgi-to-PM trafficking (17). By employing reporters with mutated basolateral PM targeting signals, they found that artificially introducing N-glycans targeted Golgi-arrested occludin and ERGIC53 chimeras to the PM while abolishing the native N-glycosylation of Fc receptor by tunicamycin arrested its exocytic trafficking in the Golgi. However, according to our knowledge, the role of O and N-glycans on the constitutive secretory transport of Golgi is still unclear and has not been systematically studied. Here, we first found that cargos lacking the O-glycosylation can accumulate in the Golgi due to their slow Golgi export. By quantitatively measuring the Golgi residence times of various transmembrane secretory reporters, we subsequently uncovered that both O- and N-glycans can function as a generic Golgi export signal at the *trans*-Golgi for the constitutive exocytic trafficking.

## Results

### Tac lacking the luminal region (Tac-TC) accumulates in the Golgi

During our study of secretory cargos, we focused on the cell surface receptor, interleukin 2 (IL2) receptor α subunit or Tac. The expression of Tac is restricted to immune cells. Devoid of known sorting signals, it is commonly used as a type I transmembrane reporter for the study of the membrane trafficking. The luminal region of Tac comprises the IL2-binding domain (IBD) and stem region (Fig. 1A). The stem region is a short stretch of ∼30 AAs connecting the transmembrane domain and the extracellular IBD. GFP-tagged truncation clones were constructed to explore roles of different regions in Tac’s secretory trafficking (Fig. 1A). As expected, full length Tac was mainly found on the PM at the steady state, with no colocalization with the Golgi marker GM130 (Fig. 1B). However, in addition to the PM, we were surprised to find a predominant Golgi localization of Tac-TC (Fig. 1B), a truncation mutant missing the entire luminal region. Tac-STC, the IBD deletion clone, displayed an intermediate phenotype, in which roughly half of expressing cells had the same localization pattern as Tac (Fig. 1B) while the remaining showed the Golgi localization (Sup. Fig. 1A)(see below for further investigation).

**Figure 1.**
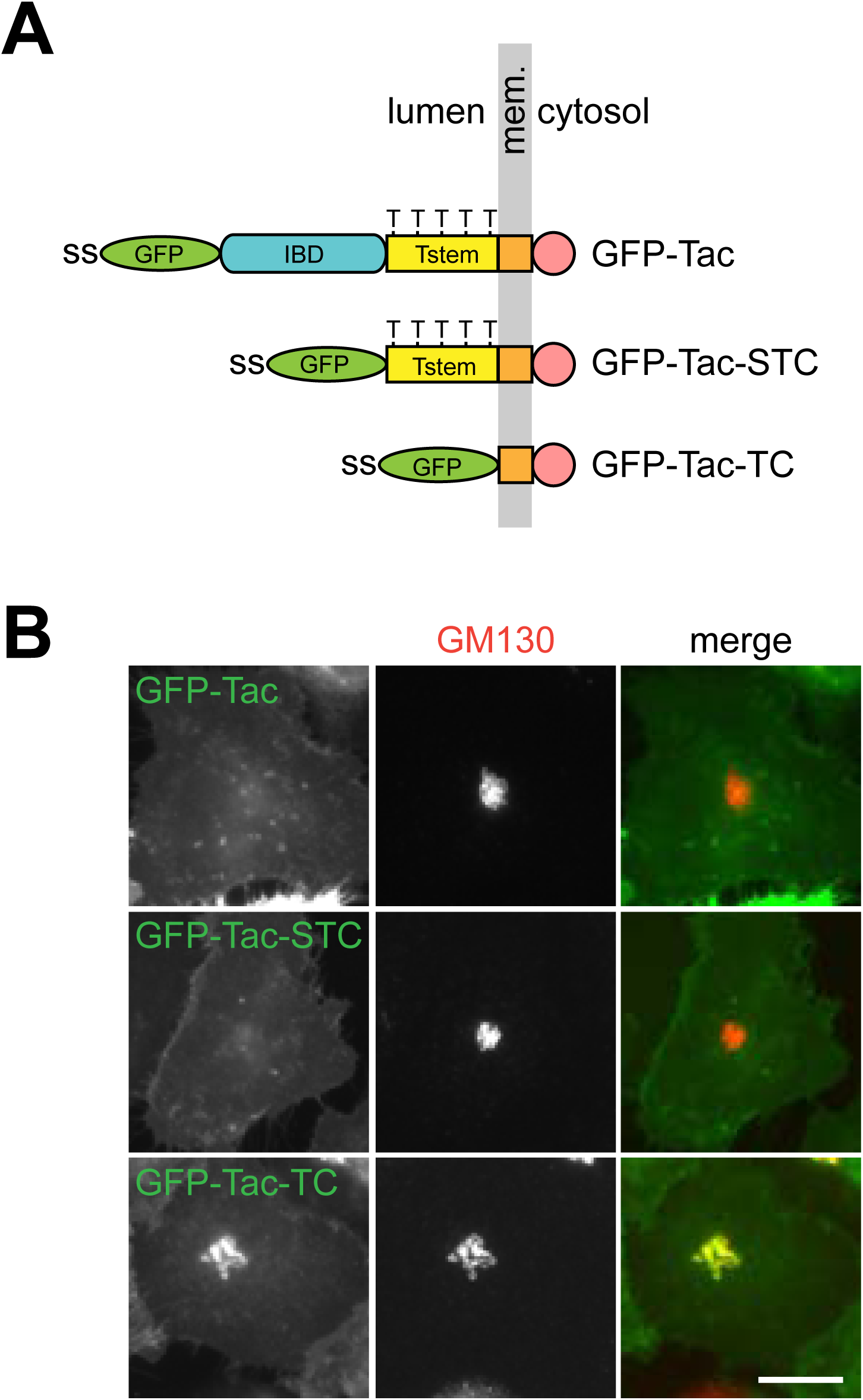
Tac-TC localizes to the Golgi. (A) The schematic diagram showing the organization of Tac truncation clones. Mem., membrane; ss, signal sequence; IBD, IL2-binding domain; Tstem, Tac stem region. Five Thr (T) residues that are potentially under mucin-type O-glycosylation are indicated in Tstem. (B) The localization of Tac truncation clones. HeLa cells transiently expressing indicated Tac truncation clone were immuno-stained for endogenous GM130 (a Golgi marker). Scale bar, 20 µm.

Similar to a conventional Golgi resident such as GalT, when the Golgi was disassembled by brefeldin A (18), the Golgi-pool of Tac-TC redistributed to the ER (Sup. Fig. 1B,C); when the Golgi was “dispersed” by the microtubule inhibitor, nocodazole (19), Tac-TC also followed GalT to the Golgi mini-stack at the ER exit site (Sup. Fig. 1D). The normal Golgi localization of Tac-TC was restored when brefeldin A or nocodazole was washed out (Sup. Fig. 1B-D). The protein synthesis inhibitor, cycloheximide (CHX), was added throughout the treatment and washout to minimize the interference of the newly synthesized Tac-TC. In summary, our observations demonstrate that Tac-TC of the Golgi-pool behaves like a conventional Golgi resident.

### The Golgi export of Tac-TC is compromised

Tac has not been known as a Golgi resident. A Golgi resident is usually expected to have a retrieval and/or retention mechanism to maintain its steady state localization at the Golgi (20,21). What is the mechanism behind the Golgi localization of Tac-TC? A secretory cargo’s Golgi localization is determined by its entrance and exit (or export) of the Golgi (Fig. 2A). For a conventional secretory cargo such as Tac, the Golgi entrance at the steady state is primarily contributed by the ER-to-Golgi transport of newly synthesized protein while the exit by the Golgi-to-PM trafficking. Although recycling can possibly occur between the Golgi and ER, e.g. the retrograde followed by the anterograde or ER-to-Golgi trafficking, the recycled fraction is likely to be small. Most importantly, if a cargo resides in the Golgi at the steady state for a sufficiently long time, the cycling between the Golgi and ER reaches a semi-steady state and hence does not produce a substantial net change in the cargo’s Golgi pool.

**Figure 2.**
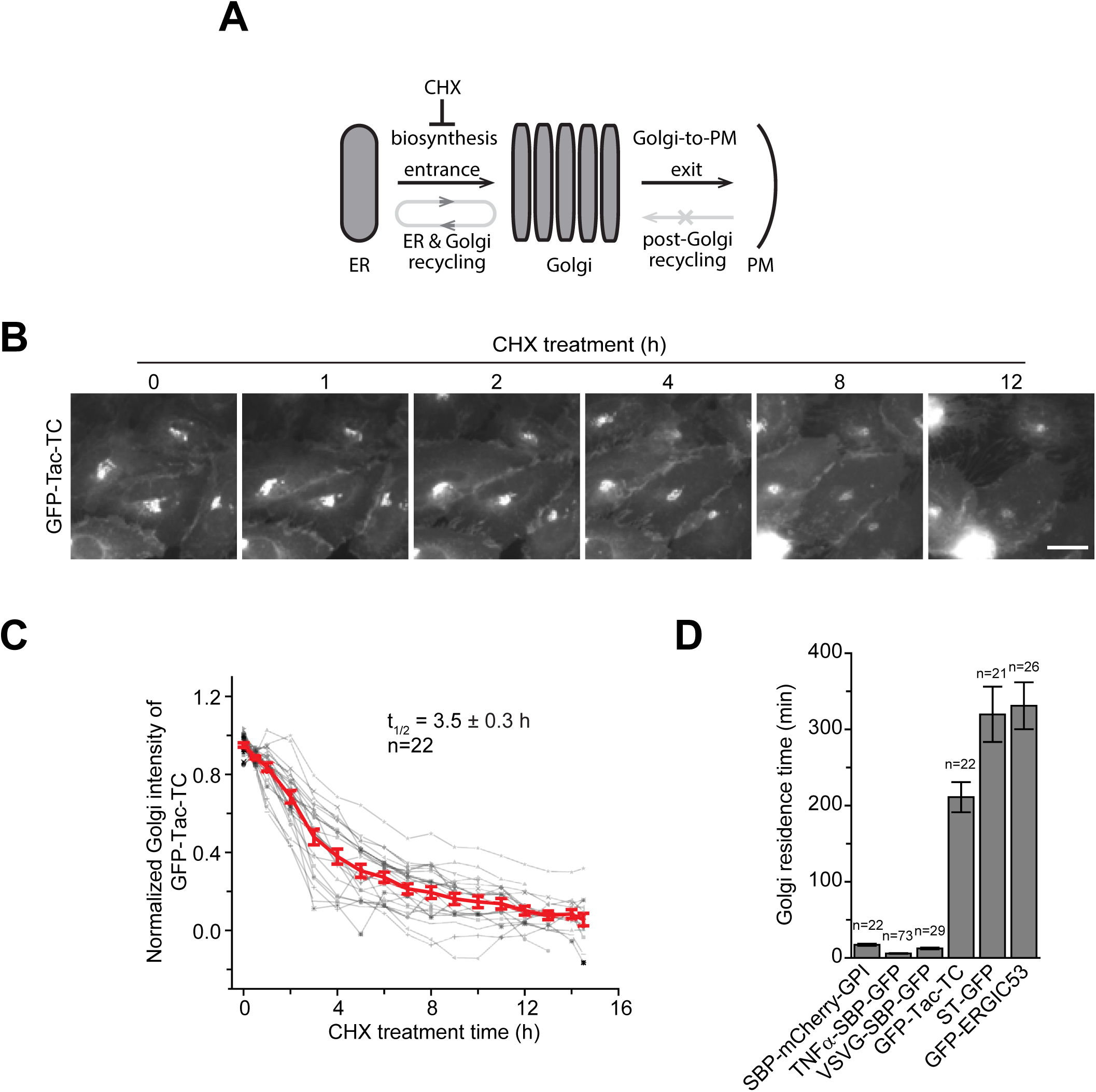
The Golgi export of Tac-TC is compromised. (A) The schematic diagram showing various trafficking pathways that contribute to a cargo’s Golgi pool. See the text for details. (B-D) The Golgi residence time of Tac-TC is significantly longer than conventional secretory cargos. In (B), HeLa cells transiently co-expressing GFP-Tac-TC and GalT-mCherry were treated with CHX and imaged live. Only GFP-Tac-TC images are shown. Scale bar, 20 µm. In (C), the total fluorescence intensity within the Golgi was quantified at each time point and subsequently plotted. Each intensity series was normalized and fitted to the first order exponential decay function to acquire the Golgi residence time, t_1/2_. Grey, individual time series; red, averaged time series. (D) The Golgi residence times of secretory cargos and Golgi residents. See Supplementary Figure 2 D-M. Error bar, mean ± stand error; n, the number of quantified cells.

We also studied the possibility of the post-Golgi retrieval of our reporters to the Golgi. It is known that VSVG does not return to the Golgi from the PM (22). Using the internalization of the surface-bound antibody, constitutive secretory cargos such as Tac, cluster of differentiation 8a (CD8a), transferrin receptor (TfR), tumor necrosis factor α (TNFα) and glycosylphosphatidylinositol anchored mCherry (mCherry-GPI), did not return to the Golgi once reaching the PM (Sup. Fig. 2A), consistent with their lack of Golgi targeting signals. Interestingly, we found that post-Golgi localized ST and ERGIC53 did not target to the Golgi from the PM and the endolysosome either (the significance of which will be discussed elsewhere) (Sup. Fig. 2B,C). Hence, the pre- and post-Golgi recycling or retrieval are not considered in the Golgi localization of cargos used in this study.

At the steady state, we can assume that the cargo’s Golgi entrance velocity is a constant, which is determined by its rate of biosynthesis, and its Golgi export velocity follows the first order kinetics. When the velocity of the export is equal to that of the entrance, the Golgi pool reaches dynamic equilibrium and its size is expected to be in a reverse relationship with the Golgi export rate constant. Hence, a cargo that slowly exits the Golgi can have a significant Golgi localization and, once the protein synthesis is blocked, the cargo’s Golgi-pool gradually decreases by following the exponential decay function (7,23).

To test the hypothesis that the Golgi localization of Tac-TC is due to its slow Golgi export, we measured the t_1/2_ of Tac-TC’s Golgi export and compared it to those of Golgi residents and conventional secretory cargos. To that end, we live-imaged cells expressing GFP-tagged proteins under the treatment of CHX (Fig. 2B). The t_1/2_, referred to as the Golgi residence time hereafter, was subsequently acquired by fitting the Golgi intensity decay data using the first order exponential function (Fig. 2C). The usage of the Golgi residence time as a Golgi export metric is advantageous as it should be independent of the cargo’s expression level and post-Golgi fates, such as degradation and cleavage. The Golgi residence times of ST and ERGIC53 were similarly measured (Sup. Fig. 2D,E). RUSH (retentioinhin using selective hooks) reporters (24) were employed to measure the Golgi residence times of constitutive secretory cargos, including mCherry-GPI, TNFα and VSVG. They were first accumulated in the Golgi by 20 °C temperature block (25) and the resulted Golgi pools were subsequently live-imaged at 37 °C (Sup. Fig. 2F-H), at which RUSH reporters started synchronously leaving the Golgi for the PM. Altogether, our quantitative data demonstrate that, though significantly less than those of ST (5.3 h) and ERGIC53 (5.5 h) (Sup. Fig. 2I,J), the Golgi residence time of Tac-TC (3.5 h) is ≥ 10-fold those of mCherry-GPI, TNFα and VSVG (Fig. 2D, Sup. Fig. 2K-M). Therefore, Tac-TC differs from conventional secretory cargos in its slow Golgi export, which likely results in its apparent Golgi localization.

### The O-glycan is essential for the efficient Golgi export of Tac

From the Golgi localization results of Tac truncation clones (Fig. 1B), we reasoned that the stem region should be essential for the efficient Golgi export. The most significant feature within this region is the presence of multiple Thr residues, which undergo mucin-type O-glycosylation in the Golgi (26). We hence asked if the O-glycan can serve as a Golgi export signal for Tac.

To test this hypothesis, we inhibited the O-glycosylation of Tac using two approaches. First, an O-glycosylation negative Tac mutant, Tac(5A), was generated by mutating all 5 Thr residues within the stem region (Fig. 3A), which were previously known to be O-glycosylated (26). When expressed in cells, Tac(5A) displayed a strong Golgi localization (Fig. 3B, 0 min), in contrast to Tac (Fig. 1B). To measure the Golgi residence time of Tac, which does not display a steady state Golgi localization, the 20 °C synchronization protocol was used to first accumulate GFP-Tac in the Golgi before live-imaging at 37 °C in the presence of CHX (Fig. 3C). Our data showed that the Golgi residence time of Tac(5A) (47 min) is almost 3-fold that of Tac (16 min) (Fig. 3D,E).

**Figure 3.**
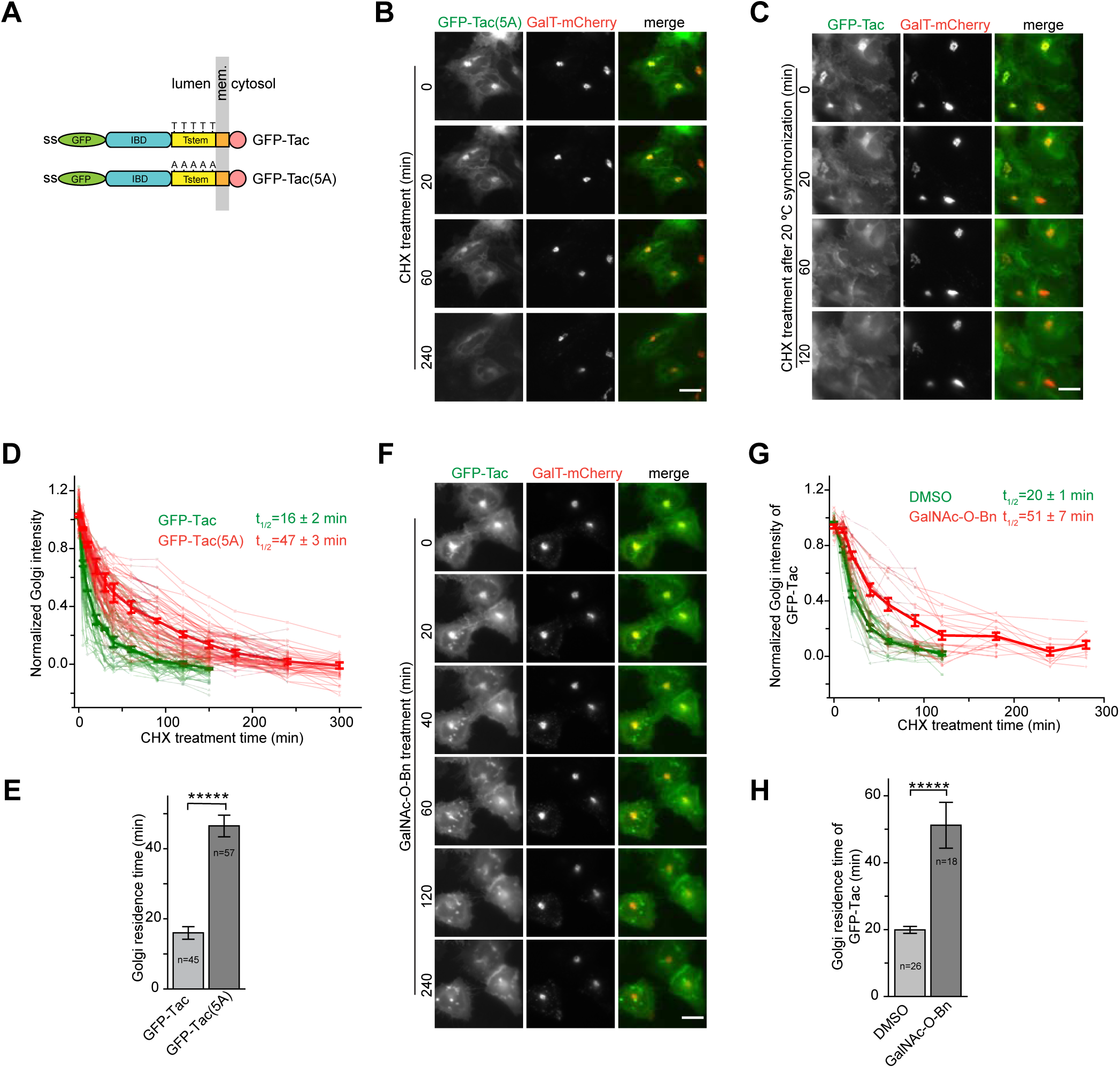
The O-glycan at the stem region is essential for the efficient Golgi export of Tac. HeLa cells were used. (A) The schematic diagram showing the domain organization and glycosylation or mutation sites of GFP-Tac and GFP-Tac(5A). The panel is organized as described in Figure 1A. Five Thr (T) residues and their corresponding Ala (A) mutations are indicated in Tstem of GFP-Tac and GFP-Tac(5A) respectively. (B-E) The Golgi residence time of Tac(5A) is significantly longer than Tac. In (B), cells transiently co-expressing GFP-Tac(5A) and GalT-mCherry were imaged live in the presence of CHX. In (C), cells transiently co-expressing GFP-Tac and GalT-mCherry were incubated at 20 °C for 4 h and subsequently warmed up to 37 °C and imaged live in the presence of CHX (see Experimental Procedures). Images acquired in (B,C) were quantified and plotted in (D,E) similar to Figure 2C,D. (F-H) Inhibiting the O-glycosylation significantly prolongs the Golgi residence time of Tac. In (F), cells transiently co-expressing GFP-Tac and GalT-mCherry were first incubated with GalNAc-O-Bn for 20 h and subsequently imaged live in the presence of CHX. Images acquired in (F) were quantified and plotted in (G) and (H) similar to Figure 2C,D. Scale bar, 20 µm; error bar, mean ± standard error; *P* values are from *t* test (unpaired and two-tailed); *****, *P* ≤ 0.000005; n, the number of quantified cells.

Second, we took advantage of a small molecule inhibitor, Benzyl 2-acetamido-2-deoxy-α-D-galactopyranoside (GalNAc-O-Bn), which compromises the mucin-type O-glycosylation by blocking the extension of O-GalNAc, the first sugar that is covalently linked to the protein (27).We confirmed the inhibitory effect of GalNAc-O-Bn on the O-glycosylation of Tac by its gel migration profile (Sup. Fig. 3B). When cells expressing GFP-Tac were treated with GalNAc-O-Bn, Tac exhibited a substantial Golgi localization and its Golgi residence time (51 min) is 2.6-fold that of dimethyl sulfoxide (DMSO) control (20 min) (Fig. 3F-H), which was acquired by the 20 °C synchronization protocol described above. Thus, extended O-glcyans (hereafter O-glycans) can substantially promote the Golgi export of Tac in comparison to O-GalNAc, which is resulted by prolonged GalNAc-O-Bn treatment. Consistent with our result, it was previously reported that O-glycosylation deficient Tac accumulated intracellularly and had little cell surface expression (28). We further investigated why the stem region containing Tac-STC displayed a mixed phenotype with a substantial fraction of cells showing the Golgi localization (Fig. 1B). The gel migration profile of Tac-STC and its 5A mutant revealed that only a small fraction of Tac-STC is O-glycosylated (Sup. Fig. 3B). Hence, its Golgi localization is probably due to the lack of O-glycosylation in Tac-STC. Together, our observations suggest that the O-glycan at the stem region might be necessary for the efficient Golgi export of Tac.

### The O-glycan might be essential for the efficient Golgi export of generic cargos

We tested two more secretory cargos that possess mucin-type O-glycosylation. Resembling Tac, CD8a is a type-I transmembrane protein that has a juxtamembrane stem region with multiple O-glycosylation sites (Fig. 4A)(29). TfR is a type-II transmembrane protein with a single O-glycosylation site (Thr104) near its transmembrane domain (Fig. 4A). VSVG, a type-I transmembrane protein that only undergoes N-glycosylation (30), served as a negative control. In cells expressing GFP-CD8a, TfR-GFP or VSVG-SBP-GFP (in the presence of biotin), we observed that, while none appeared at the Golgi in control treatment, CD8a and TfR, but not VSVG, displayed strong Golgi localization under GalNAc-O-Bn treatment (Fig. 4B,C; Sup. Fig. 4A). Their Golgi residence times under control and GalNAc-O-Bn treatment were subsequently measured (Fig. 4D-J). We found that GalNAc-O-Bn treatment substantially prolonged the Golgi residence times of both CD8a and TfR, but not that of VSVG, therefore suggesting that the O-glycan might be necessary for the efficient Golgi export of generic O-glycosylated cargos.

**Figure 4.**
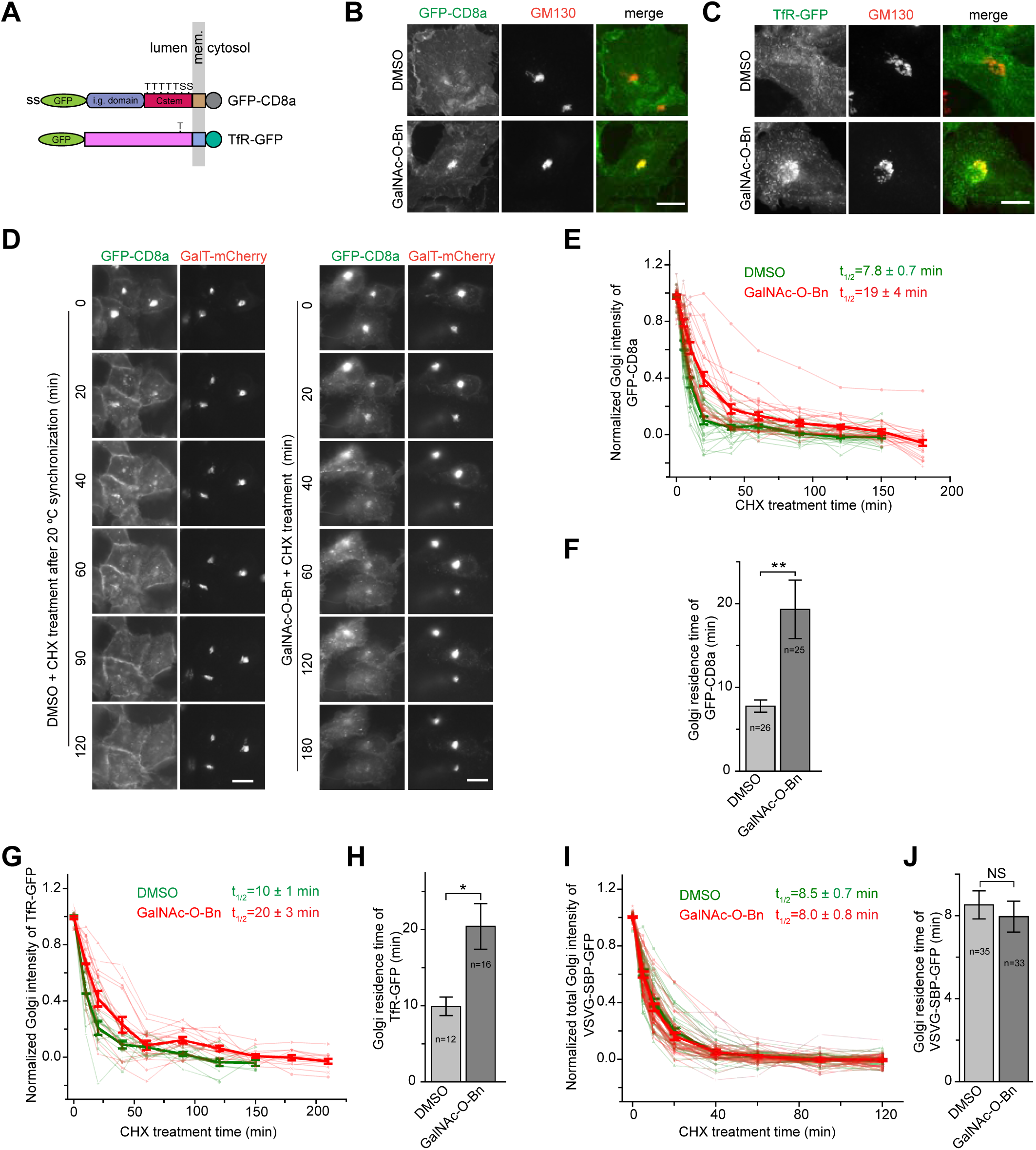
The O-glycan is required for the efficient Golgi export of CD8a and TfR. HeLa cells were used. (A) The schematic diagram showing the domain organization and potential O-glycosylation sites of CD8a and TfR. The panel is organized as described in Figure 1A. i.g., immunoglobulin. Thr (T) or Ser (S) residues that are potentially under mucin-type O-glycosylation are indicated. (B,C) A substantial amount of CD8a and TfR is accumulated in the Golgi when their O-glycosylation is inhibited. Cells transiently expressing GFP-CD8a or TfR-GFP were treated with DMSO or GalNAc-O-Bn for 20 h before immuno-staining of endogenous GM130. (D-J) Inhibiting the O-glycosylation significantly prolongs the Golgi residence times of CD8a and TfR but not VSVG. In (D), cells transiently co-expressing GFP-CD8a and GalT-mCherry were treated with either DMSO or GalNAc-O-Bn for 20 h. Both DMSO and GalNAc-O-Bn treated cells were subsequently incubated at 20 °C to accumulate GFP-CD8a in the Golgi before time-lapse imaging at 37 °C in the presence of CHX (see Materials and methods). Images acquired from (D) were quantified and plotted in (E,F) as Figure 2C,D. In (G,H), the Golgi residence time of TfR under DMSO or GalNAc-O-Bn treatment was similarly acquired and plotted as in (E,F). See Supplementary Figure 4B,C for images. In (I, J), after the treatment of DMSO or GalNAc-O-Bn for 20 h, cells transiently co-expressing VSVG-SBP-GFP and GalT-mCherry were treated with biotin at 20 °C to accumulate VSVG in the Golgi. Time-lapse imaging was subsequently conducted in the presence of biotin and CHX at 37 °C and the Golgi residence time was acquired and plotted as (E,F). See Supplementary Figure 4D,E for images. Scale bar, 20 µm; error bar, mean ± standard error; *P* values are from *t* test (unpaired and two-tailed); *, *P* ≤ 0.05; **, *P* ≤ 0.005; NS, not significant; n, the number of quantified cells.

### The O-glycan is sufficient to promote the Golgi export of Tac(5A)

We investigated the Golgi export of GFP-Tac(5A) (Fig. 3A) when O-glycans are re-introduced. The stem region of CD8a or Cstem (AA 134-177) has 5 Thr and 2 Ser for potential O-glycosylation (Fig. 5A; Sup. Fig. 5A) (29). Cstem was adopted as a transferable sequence for multiple O-glycosylation, together with a non-glycosylation negative control, Cstem(7A), in which the above mentioned Thr and Ser are mutated to Ala. Cstem or Cstem(7A) was subsequently inserted in between GFP and Tac(5A) to make chimeras, GFP-Cstem-Tac(5A) and GFP-Cstem(7A)-Tac(5A) (Fig. 5A; Sup. Fig. 5A). Both chimeras were likely N-glycosylated as their gel migration profiles were sensitive to PNGase F treatment (Sup. Fig. 5B). Furthermore, GFP-Cstem(7A)-Tac(5A) was observed to migrate slightly faster than GFP-Cstem-Tac(5A), consistent with the expectation that the former has less or no O-glycans. When their Golgi residence times were subsequently measured, we observed that GFP-Tac(5A) with Cstem (38 min) had a much-reduced Golgi residence time in comparison to that with Cstem(7A) (76 min) (Fig. 5B,C; Sup. Fig. 5C), suggesting that the O-glycan is probably sufficient to promote the Golgi export of Tac(5A). Altogether, our data demonstrate that the O-glycan is both necessary and sufficient for the efficient Golgi export of Tac, therefore supporting our hypothesis that the O-glycan might function as a Golgi export signal.

**Figure 5.**
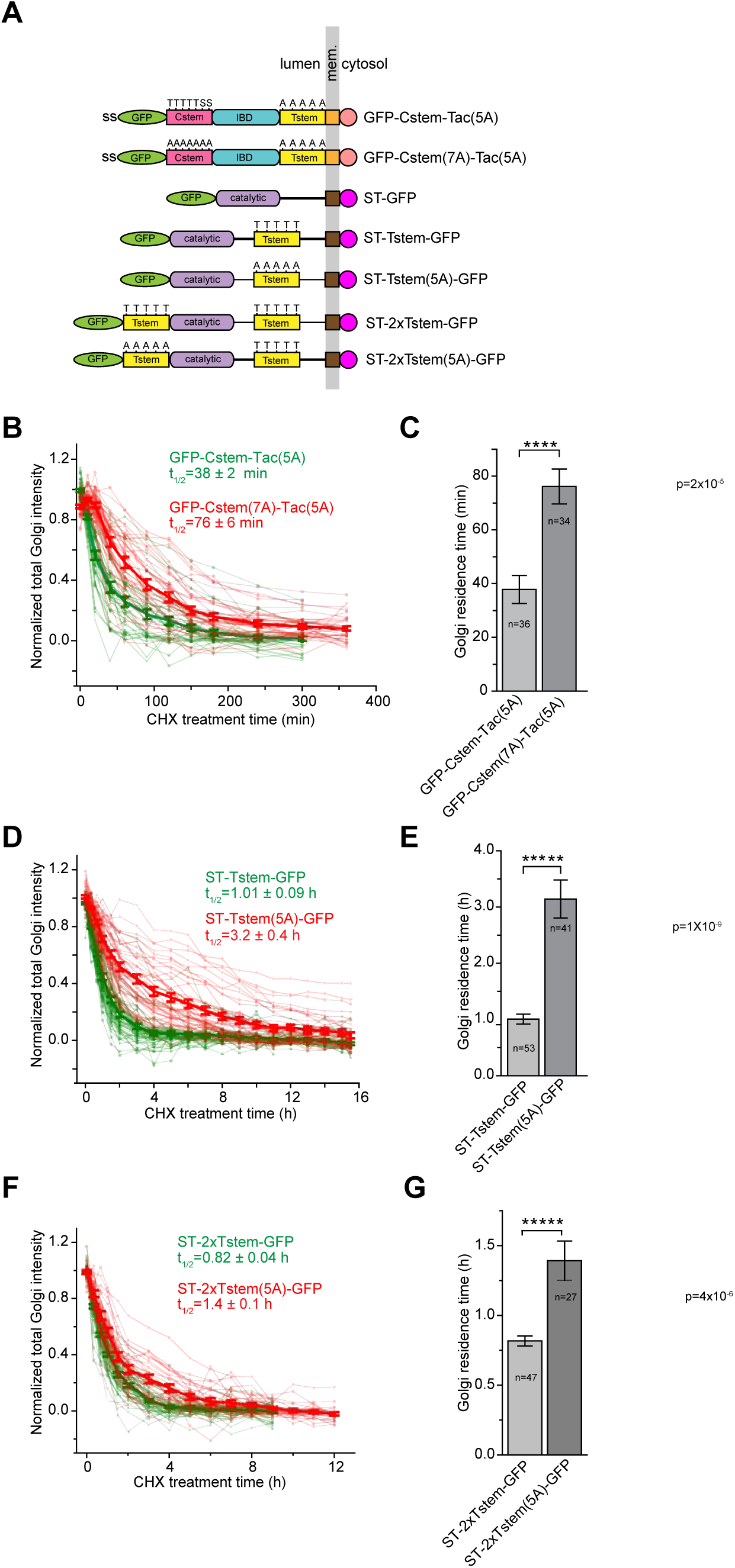
The O-glycan can promote the Golgi export in an additive manner. (A) The schematic diagram showing the domain organization of Tac and ST chimeras. The panel is organized as described in Figure 1A. Thr (T) and Ser (S) residues that are potentially O-glycosylated together with corresponding Ala (A) mutations are indicated above Cstem or Tstem. Catalytic, catalytic domain of ST. (B-G) The Golgi residence times were acquired and plotted similar to Figure 2B,C. (B,C) demonstrate that introducing the O-glycan reduces the Golgi residence time of Tac(5A), therefore suggesting that the O-glycan is sufficient to promote the Golgi export. The Golgi residences times shown in (D,E) indicate that Tstem in ST-Tstem-GFP are likely O-glycosylated. (F,G) demonstrate that extra O-glycans, introduced by the membrane distal Tstem, promote the efficient Golgi export of ST-2xTstem-GFP in comparison with ST-2xTstem(5A)-GFP. Error bar, mean ± standard error; *P* values are from *t* test (unpaired and two-tailed); ****, *P* ≤ 0.00005; *****, *P* ≤ 0.000005; n, the number of quantified cells.

### The O-glycan promotes the Golgi export in an additive manner

We chose ST to study the effect of O-glycan’s quantity on the Golgi export. We first confirmed that ST is N-glycosylated by PNGase F digestion (Sup. Fig. 5D), as previously reported (31). Although ST is predicted to be O-glycosylated in its juxtamembrane sequence (32), we further introduced Tac stem region (Tstem) (Sup. Fig. 5A), a known multi-O-glycosylation sequence (26), to ST. Tstem or its 5A mutant, Tstem(5A), was inserted to ST’s juxtamembrane sequence to make ST-Tstem-GFP or ST-Tstem(5A)-GFP respectively (Fig. 5A; Sup. Fig. 5E). In these chimeras, the added O-glycosylation or mutant sites are proximal to the membrane. Both chimeras were N-glycosylated as their gel migration profiles were sensitive to PNGase F treatment and ST-Tstem(5A)-GFP always migrated slightly faster than ST-Tstem-GFP (Sup. Fig. 5F), consistent with the prediction that the latter should have more O-glycans. When their Golgi residence times were subsequently measured, ST-Tstem-GFP displayed a significantly shorter Golgi residence time (1.01 h) than ST-Tstem(5A)-GFP (3.2 h) (Fig. 5D,E), further supporting our finding that O-glycosylation promotes the Golgi export.

To introduce more O-glycans, an extra copy of Tstem or Tstem(5A), which serves as a negative control, was inserted in between GFP and the catalytic domain of ST to make chimera ST-2xTstem-GFP or ST-2xTstem(5A)-GFP respectively (Fig. 5A; Sup. Fig. 5E). In these two chimeras, the newly added O-glycosylation and mutant sequences are distal from the membrane. During the subsequent gel analysis, we observed that ST-2xTstem-GFP migrated slower than ST-2xTstem(5A)-GFP (Sup. Fig. 5G), consistent with the prediction that the former has more O-glycosylation sites than the latter. We subsequently measured that the Golgi residence time of ST-2xTstem-GFP (0.82 h) was significantly shorter than that of ST-2xTstem(5A)-GFP (1.4 h). In summary, our observation that a cargo with more O-glycans can have a shorter Golgi residence time implies that the O-glycan might promote the Golgi export in an additive manner.

### The N-glycan might also function as a Golgi export signal

We asked whether the N-glycan can also function as a Golgi export signal. While the O-glycan is mostly found in the unfolded or disordered regions, such as inter-domain linkers or loops, the N-glycosylation mostly occurs in protein domains and is essential for the proper folding of a protein in the ER (32). To circumvent the requirement of the N-glycosylation in the folding and subsequent ER export of cargos, we chose the N-glycosylated GFP since GFP can fold in the ER without the requirement of glycosylation. Missing the luminal region of Tac, GFP-Tac-TC does not undergo N-glycosylation. A sequon N-glycosylation site was introduced to a loop connecting β6 and β7 of GFP to make GFP(N-glyc) (33) and GFP(N-glyc)-tagged Tac-TC was constructed (Fig. 6A). In the subsequent gel analysis, we observed that GFP(N-glyc)-Tac-TC migrated slower than GFP-Tac-TC and, after PNGase F treatment, both chimeras migrated at the same speed (Sup. Fig. 6), demonstrating the successful N-glycosylation of GFP(N-glyc)-Tac-TC. Our live-imaging data showed that the Golgi residence time of GFP(N-glyc)-Tac-TC (1.2 h) was reduced to almost one third that of GFP-Tac-TC (3.5 h) (Fig. 6B-D), therefore suggesting that the N-glycan might function as a Golgi export signal, similar to what we found for the O-glycan.

**Figure 6.**
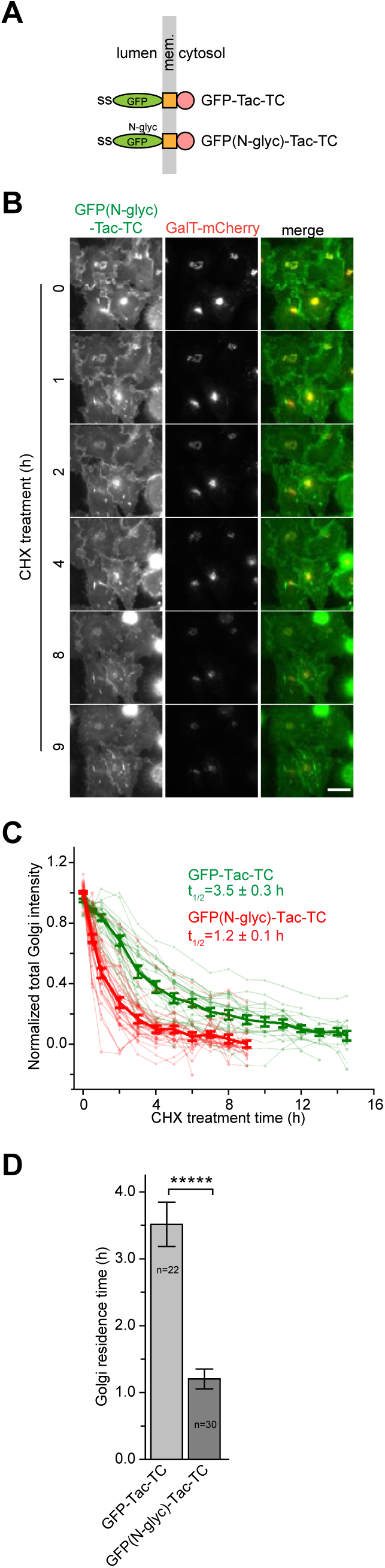
The N-glycan can accelerate the Golgi export of GFP-Tac-TC. HeLa cells were used. (A) The schematic diagram showing the domain organization of Tac-TC fused to N-glycosylated (N-glyc) or wild type GFP. The panel is organized as described in Figure 1A. (B-D) The introduction of the N-glycan significantly promotes the Golgi export of GFP-Tac-TC. The experiment and panel organization are similar to those of Figure 2B-D. GFP-Tac-TC data are from Figure 2C. Scale bar, 20 µm; error bar, mean ± standard error; *P* values are from *t* test (unpaired and two-tailed); *****, *P* ≤ 0.000005; n, the number of quantified cells.

### Tac-TC accumulates at the interior of the *trans*-Golgi cisternae

To compare the intra-Golgi transport of Tac-TC with Tac, corresponding RUSH-reporters were constructed (Fig. 7A). During the chase, the synchronized trafficking of Tac-TC and Tac was quantitatively monitored by the localization quotient (LQ), a linear metric of the axial Golgi localization using our recently developed Golgi super-resolution tool, the Golgi protein localization by imaging centers of mass (GLIM) (34). Both Tac-TC and Tac rapidly entered the secretory pathway from the ER and transited from the *cis* to *trans*-side of the Golgi, as shown by increases of their LQs (Fig. 7B,C). t_1/2_s from the first order exponential fitting of the LQ vs time plots reflect the intra-Golgi transport velocity. Similar to their Golgi export, the intra-Golgi transport velocity of Tac-TC (t_1/2_= 17.1 min) is slower than that of Tac (t_1/2_= 5.9 min). Like Tac (Fig. 7B) and other constitutive secretory cargos that we previously studied (34), the LQ of Tac-TC plateaued at ∼ 1 (Fig. 7C), a localization roughly corresponding to the *trans*-Golgi (34). Consistently, steady state LQs of Tac-TC and Tac(5A), which were acquired from cells expressing respective GFP-tagged construct, were measured to be 0.94 ± 0.02 (n=118) and 0.95 ± 0.02 (n=123). The observation that the slow Golgi exit cargo accumulates at the *trans*-Golgi provides further evidence that the *trans*-Golgi instead of the TGN is the export site of constitutive secretory cargos. By identifying *en face* views of nocodazole-induced Golgi mini-stacks (35), we found that the *trans*-localized Tac-TC and Tac(5A) mainly distribute to the interior of Golgi cisternae under Airyscan super-resolution microscopy (Fig. 7D-F and Sup. Fig. 7), similar to other secretory cargos that we previously studied (35).

**Figure 7.**
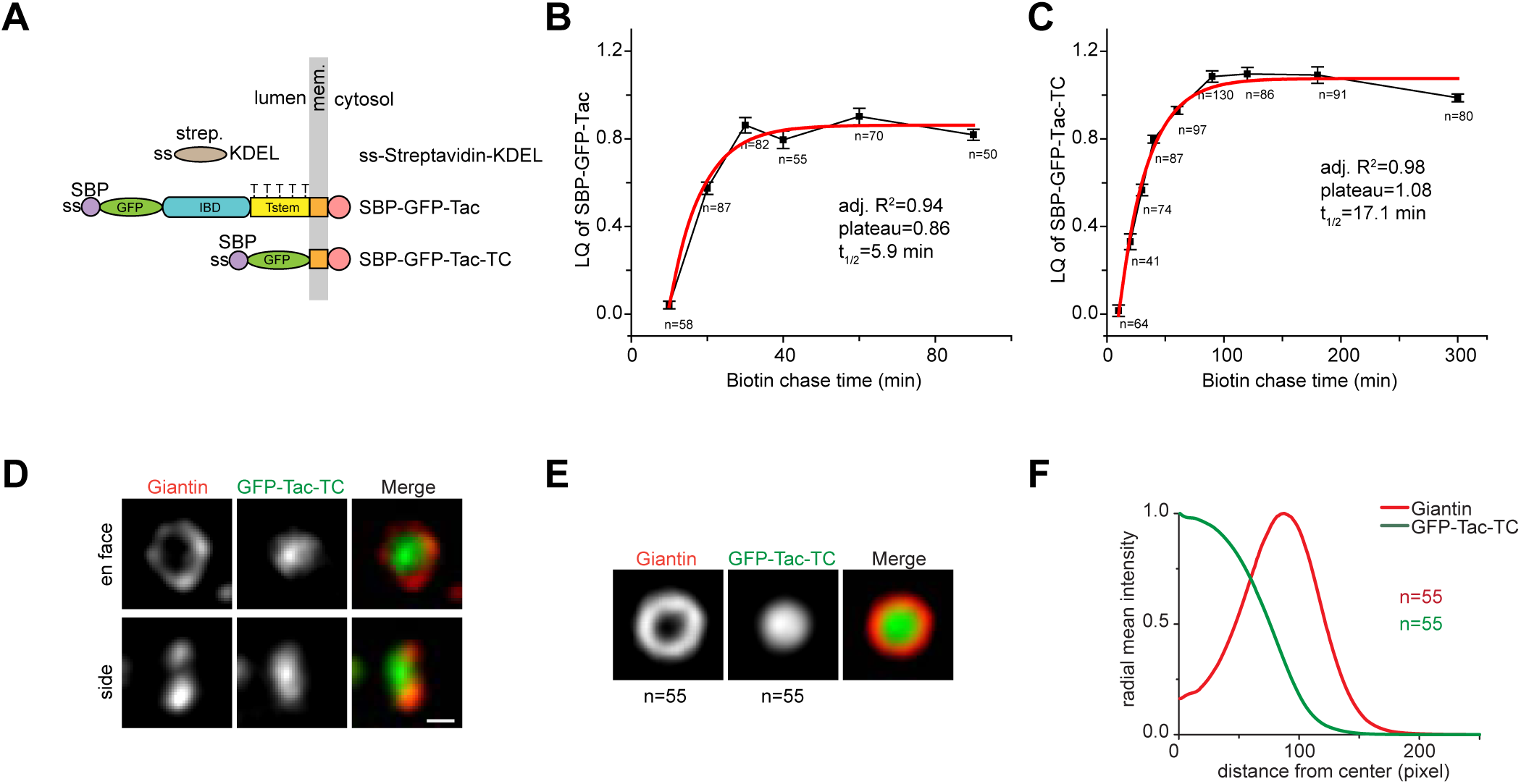
Tac-TC accumulates at the interior of the *trans*-Golgi cisternae. HeLa cells were used. (A) The schematic diagram showing the domain organization of Tac and Tac-TC RUSH reporters. The panel is organized as described in Figure 1A. ss-Streptavidin-KDEL is the ER hook. Strep., streptavidin; SBP, streptavidin-binding peptide. The five Thr residues that are potentially under mucin-type O-glycosylation are indicated on Tstem. (B,C) The intra-Golgi transport kinetics of Tac and Tac-TC as revealed by the LQ vs time plot. Cells transiently co-expressing the RUSH reporter, SBP-GFP-Tac or SBP-GFP-Tac-TC, and GalT-mCherry were treated with nocodazole to induce the formation of Golgi ministacks. The ER arrested RUSH reporter was subsequently released by the administration of biotin. At different time points, cells were immuno-stained for endogenous GM130 and the LQs for the RUSH reporter were acquired and fitted to the first order exponential function. Error bar, mean ± standard error; n, the number of Golgi ministacks quantified. Adj. R^2^, adjusted R^2^. (D-F) Tac-TC localizes to the interior of Golgi cisternae at the steady state. (D) shows Airyscan super-resolution images of the en face and side view of Giantin and GFP-Tac-TC in the Golgi ministack. Cells transiently expressing GFP-Tac-TC were treated with nocodazole to induce the formation of Golgi ministacks and stained for endogenous Giantin. The en face and side view images were selected. Scale bar, 500 nm. The en face averaging of Giantin and GFP-Tac-TC in the Golgi ministack is shown in (E). The corresponding radial mean intensity profile is displayed in (F) with the distance from the center of fluorescence mass (Giantin peak is normalized to 100) as the x-axis and the radial mean intensity (normalized) as the y-axis. n, the number of Golgi ministack images used.

## Discussion

The role of glycans in the secretory trafficking is still unclear. Here, by quantitatively measuring the Golgi residence times of transmembrane secretory reporters, we uncovered that both O and N-glycans can function as a signal for the efficient Golgi export. Abolishing or inhibiting the O-glycosylation of Tac, CD8a and TfR by mutagenesis or small molecule substantially slowed down their Golgi export and resulted in their Golgi localization or retention, while adding O-glycans to Golgi-localized Tac(5A) and ST produced the opposite effect, that is the promotion of their Golgi export and the reduction of their Golgi retention. Similarly, introducing the N-glycan to GFP also accelerated the Golgi export of the engineered reporter, GFP-Tac-TC. Using the conventional Golgi resident ST as a reporter, we demonstrated that the Golgi export promotion effect of the O-glycan can be additive. Therefore, glycans can be necessary and sufficient for the efficient Golgi export and our discovery might be extended to generic transmembrane proteins, including conventional Golgi residents and transiting secretory cargos.

It has been previously reported that the apical targeting of cargos is compromised when their N- or O-glycosylation was disrupted (15,16). Retrospectively and considering our current finding, the compromised apical targeting is likely due to the slow Golgi export of glycan-deficient cargos. The O-glycosylation at the stem region has also been documented to protect cell surface proteins such as low-density lipoprotein receptor (36), TfR (37), decay accelerating factor (38) and Tac (39) from ectodomain shedding. Therefore, our discovery further expands the role of the O-glycan on the expression of cell surface receptors by suggesting that it contributes to their Golgi-to-PM or exocytic trafficking as well.

Most protein trafficking signals are cytosolic AA sequences, which are recognized by trafficking machinery components such as vesicular coats and adaptors. Hence, as a Golgi export signal, the glycan is unique since it is a non-proteinaceous moiety and confined to the lumen. Probably, it makes the biological sense for the glycan to function as a Golgi export signal as it ensures enough glycosylation of secretory cargos before leaving the Golgi. What type of sorting information does the glycan encode? And how is it recognized and recruited to an exocytic carrier by the trafficking machinery? It has been previously postulated that glycans might interact with lipid rafts or lectins such as VIP36 for their roles in the apical targeting (15,16). We argue that, different from conventional trafficking signals, glycans except M6P (11,12) are unlikely to be sorted by a receptor-mediated mechanism due to their vast diversity in structure and composition. Instead, we speculate that their sorting is primarily based on physico-chemical properties of the glycan such as the size, shape and hydrophilicity.

Although other mechanisms are certainly possible, our data are compatible with a simple model in which the sorting of glycans to exocytic carriers is by the molecular exclusion mechanism, which is elaborated below. Previous imaging data have demonstrated that the interior of the medial and *trans*-Golgi cisternae is tightly packed with arrays of Golgi glycosylation enzymes (40,41), which constitute what we termed “the Golgi enzyme matrix”. During the cisternal transition of a secretory cargo from the *cis* to *trans*-side of the Golgi, the cargo mainly partitions to the Golgi enzyme matrix (35). In this process, the addition of glycans increases the hydrodynamic volume or hydrophilicity of the cargo’s luminal region, probably resulting in its exclusion from the enzyme matrix. Hence, the expelled but fully glycosylated cargo ends up in the rim of Golgi cisternae, where trafficking machinery components subsequently pack it into the exocytic carrier (35). Using the analogy of the gel filtration column, a non- or under-glycosylated cargo probably has a small Stokes radius and therefore a large elution volume or long Golgi residence time, while the glycosylated cargo has the opposite as a result of its increased size. Specially, for GPI-anchored proteins, the juxtamembrane glycan moiety might directly serve as a Golgi export signal. For the non-glycosylated cargos, which are of minority, other mechanisms are probably utilized to facilitate their efficient Golgi export. For example, multimerization may increase the hydrodynamic volume so that cargos are excluded from the interior of the Golgi enzyme matrix. Indeed, we previously demonstrated that large soluble secretory proteins always localize to the rim of Golgi cisternae during their intra-Golgi transport (35). Further experiments are certainly required to test this view of the Golgi export.

## Experimental Procedures

### DNA plasmids

See Supplementary Table 1.

### Antibodies, enzyme and small molecules

Mouse anti-Lamp1 monoclonal antibody (mAb) (H4A3; 1:500 for immunofluorescence or hereafter IF), mouse anti-CD63 mAb (H5C6; 1:100 for IF), and mouse anti-CD8a mAb (OKT8; 1:500 for IF) were from Developmental Studies Hybridoma Bank. Mouse anti-GFP mAb (#sc-9996; 1:1000 for Western blot) was purchased from Santa Cruz. Mouse anti-EEA1 mAb (#610456; 1:500 for IF) and mouse anti-GM130 mAb (#610823; 1:500 for IF) were from BD Biosciences. Horseradish peroxidase (HRP)-conjugated goat anti-mouse (#176516; 1:10,000 for Western blot) was from Bio-Rad. Alexa Fluor-conjugated goat antibodies against mouse IgG (1:500 for IF) were from Thermo Fisher Scientific.

PNGase F was purchased from New England Biolab. The following small molecules were commercially available: nocodazole (Merck; working concentration: 33 µM), brefeldin A (Epicenter Technologies; working concentration: 10 µg/ml), cycloheximide (Sigma Aldrich; working concentration: 10 µg/ml), GalNAc-O-Bn (Sigma Aldrich; working concentration: 2 mM).

### Production of rabbit anti-mCherry polyclonal antibody

The plasmid DNA encoding His-mCherry (mCherry-pET30ax) was used to transform BL21DE3 *E*.*coli* cells and resulting bacteria were induced by 0.25 mM isopropyl β-D-thiogalactopyranoside (IPTG) (Thermo Fischer Scientific) at 16°C overnight. Bacterial cells were pelleted by centrifugation and lysed by sonication in phosphate buffered saline (PBS) with 8 M Urea. The supernatant was collected by centrifugation and incubated with nickel-nitrilotriacetic acid agarose beads at room temperature for 2 h. The beads were washed by PBS supplemented with 8 M urea and 20 mM imidazole and eluted using PBS supplemented with 8 M urea and 250 mM imidazole. The eluted His-mCherry was concentrated and the buffer was changed to PBS containing 4 M urea. His-mCherrry was used as an antigen to immunize rabbits and anti-sera were collected by Genemed Synthesis Inc.

The plasmid DNA encoding GST-mCherry (mCherry-pGEB) was used to transform BL21DE3 *E*.*coli* cells. Bacterial cells were induced by 0.25 mM IPTG at 16°C overnight and collected by centrifugation, followed by sonication in the lysis buffer containing 50 mM Tris pH 8.0, 100 mM NaCl, 0.1% Triton X-100 and 1 mM Dithiothreitol (DTT, Sigma-Aldrich). After centrifugation, the collected supernatant was incubated with Glutathione Sepharose beads (GE Healthcare) at 4 °C overnight and washed by the lysis buffer. After further washing by 200 mM sodium borate buffer (pH 9.0), GST-mCherry-immobilized on beads was cross-linked to beads by incubating with sodium borate buffer supplemented with 50 mM dimethyl pimelimidate (Sigma-Aldrich), followed by neutralization with 200 mM ethanolamine (pH 8.0). Cross-linked beads were incubated with mCherry anti-sera for 1 h at room temperature. After PBS washing, the bound mCherry antibody was eluted by 100 mM glycine pH 2.8, neutralized by 1 M Tris pH 8.0, and dialyzed in PBS.

### Cell culture and transfection

HeLa cells were cultured in Dulbecco’s Modified Eagle’s Medium supplemented with 10% fetal bovine serum (FBS). For IF labeling, cells were seeded onto a F 12 mm glass coverslip in a 24-well plate. For fluorescence live-imaging, cells were seeded onto a F 35 mm glass-bottom Petri-dish (Cellvis) and imaged in the CO_2_-independent medium (Thermo Fisher Scientific) supplemented with 4 mM glutamine and 10 % FBS at 37 °C. Lipofectamine 2000 (Thermo Fisher Scientific) was used to transfect cells according to standard protocol from the manufacturer. Experiments were conducted ∼ 20 h after transfection.

### IF labeling

Cells grown on F 12 mm glass coverslips were fixed by 4% paraformaldehyde in PBS. After washing, cells were incubated with 100 mM NH_4_Cl to block the residual paraformaldehyde and subjected to further wash in PBS. Next, cells were sequentially incubated with the primary antibody followed by the secondary antibody (Alexa Fluor-conjugated goat anti-mouse and/or rabbit antibodies) diluted in PBS containing 5% FBS, 2% bovine serum albumin and 0.1% saponin (Sigma-Aldrich). A washing step was included between the primary and secondary antibody incubation. After extensive wash in PBS, labelled cells were mounted in Mowiol mounting medium, containing 12% Mowiol 4-88 (EMD millipore), 30% glycerol and 100 mM Tris pH 8.5. Once dried, coverslips were sealed in nail polish.

### Conventional fluorescence microscopy

Live and fixed cells were imaged under Olympus IX83 inverted wide-field microscope system equipped with an oil 100×/NA 1.40 objective (Plan Apo), an oil 63×/NA 1.40 objective (Plan Apo) and an oil 40×/NA 1.20 objective (fluorite), a motorized stage, a focus drift correction device, a 37 °C enclosed environment chamber, motorized filter cubes, a scientific complementary metal oxide semiconductor camera (Neo; Andor) and a 200 W metal-halide excitation light source (Lumen Pro 200; Prior Scientific). Filters and dichroic mirrors were optimized for GFP/Alexa Fluor 488, mCherry/Alexa Fluor 594 and Alexa Fluor 680. The microscope system was controlled by MetaMorph (Molecular Devices).

### Super-resolution fluorescence microscopy

This was conducted using Airyscan super-resolution microscope system (Carl Zeiss) as previously described (35).

### Measuring LQs

This was described previously (34,41).

### Golgi residence time quantification

HeLa cells grown on a F 35 mm glass-bottom Petri-dish were co-transfected with a fluorescence protein-tagged Golgi marker, GalT-mCherry or GFP-GM130 together with indicated GFP or mCherry-tagged reporter for about 20 h. For reporters that have substantial steady-state Golgi localization, cells were directly subjected to live-imaging in the CO_2_-independent medium (Thermo Fisher Scientific) supplemented with 10% FBS, 4 mM glutamine and 10 µg/ml CHX at 37 °C. At the steady state, GFP-Tac, GFP-CD8a and TfR-GFP do not have significant Golgi localization. To accumulate or synchronize these reporters in the Golgi, cells expressing them were incubated at 20 °C for 3 h in the absence of CHX followed by 20 °C for 1 h before live-imaging at 37 °C in the presence of CHX. For RUSH reporters, cells expressing them were first incubated with 50 ng/ml streptavidin to quench residual amount of biotin present in the tissue culture medium. After 20 h, cells were washed extensively to remove the streptavidin and incubated with 40 µM biotin and 10 µg/ml CHX at 20 °C to accumulate RUSH reporters in the Golgi before live-imaging at 37 °C.

For each reporter, 2D time-lapse imaging was conducted until the reporter fluorescence signal in the Golgi almost disappeared. Typically, < 20 frames were acquired during each time-lapse imaging and therefore the photobleaching was negligible. Golgi residence times were subsequently quantified from these imaging data using ImageJ (https://imagej.nih.gov/ij/). In a time-lapse, Golgi ROIs were generated by the intensity segmentation of co-expressed GalT-mCherry or -GFP with manual modifications. The total Golgi intensity of the reporter at each time point was measured in ImageJ and fitted by the first order exponential decay function, y=y_0_ + A_1_ exp(-(x-x_0_)/t_1_), in OriginPro8.5 (OriginLab). The Golgi residence time, which is defined as the t_1/2_, was then calculated as 0.693*t1. Only time-lapses with adjusted R^2^ ≥ 0.80 and the length of acquisition ≥ 1.33*t_1/2_ were considered in our calculation.

### PNGase F digestion

HeLa cells seeded in a 24-well plate were transiently transfected with the reporter DNA plasmid. After centrifugation, pellets were suspended in 10 µl glycoprotein denaturing buffer containing 0.5% SDS and 40 mM DTT and heated at 100 °C for 10 min to denature proteins. Next, 2 µl GlycoBuffter 2 (10x, 500 mM Sodium Phosphate, pH 7.5), 2 µl 10% NP-40, 6 µl H_2_O and 1 µl PNGase F (New England Biolab) were added into the cell lysate and the mixture was incubated at 37 °C for 1 h. After adding SDS-sample buffer, the digested cell lysate was analyzed by standard SDS-PAGE followed by Western blot.

### GalNAc-O-Bn treatment

HeLa cells transiently expressing the reporter of interest were incubated with the tissue culture medium containing 2 mM GalNAc-O-Bn or DMSO as a control for 20 h before fluorescence imaging or immuno-blotting.

### VHH (variable heavy-chain domain of heavy-chain-only antibody)-mCherry purification

6×His-tagged VHH-mCherry (Addgene #109421) was transformed into BL21DE3 *E*.*coli* cells. Bacterial culture was induced by 0.25 mM IPTG at 16°C overnight. Cells were harvested by centrifugation and resuspended using the lysis buffer (100 mM HEPES pH 7.4, 500 mM KCl, 1% Triton X-100, 200 μg/ml lysozyme, 0.5 mM phenylmethylsulfonyl fluoride, 10 mM imidazole, and 2 mM DTT), followed by sonication lysis. After centrifugation, the resulting supernatant was incubated with nickel-nitrilotriacetic acid agarose beads (QIAGEN) pre-equilibrated with the washing buffer (20 mM HEPES, pH 7.4, 200 mM KCl, 10% Glycerol, and 2 mM DTT) supplemented with 10 mM Imidazole at 4 °C overnight. Beads were then washed by the washing buffer supplemented with 25 mM imidazole and eluted by the washing buffer supplemented with 250 mM imidazole. The elute was dialyzed in the PBS at 4 °C overnight and quantified by Coomassie blue stained SDS-PAGE gel using the bovine serum albumin as standard.

### Cell internalization assay

HeLa cells transiently expressing indicated GFP-tagged protein, CD8a-furin-mEOS2 or SBP-mCherry-GPI were incubated with 5 µg/ml VHH-mCherry, mouse anti-CD8a mAb or rabbit anti-mCherry polyclonal antibody respectively for 2 h followed by fixation and immuno-staining for indicated proteins.

## Author contribution

L.L. conceived and supervised the study. L.L. and X.S. designed the experiments. X.S., H.C.T. and B.C performed experiments. X.S., H.C.T. and L.L. analyzed the data. L.L. and X.S. wrote the manuscript.

## Acknowledgement

We would like to thank the following researchers for sharing their DNA plasmids with us: F. Perez, H. Hauri, T. Kirchhausen, J. Lippincott-Schwartz, B. Gumbiner and E. Snapp. This work was supported by the following grants to L.L.: MOE AcRF Tier1 RG35/17, Tier2 MOE2015-T2-2-073 and Tier2 MOE2018-T2-2-026.

## Competing interests

The authors declare that they have no competing financial interests.

## Supplementary figure legends

**Supplementary Figure 1.**
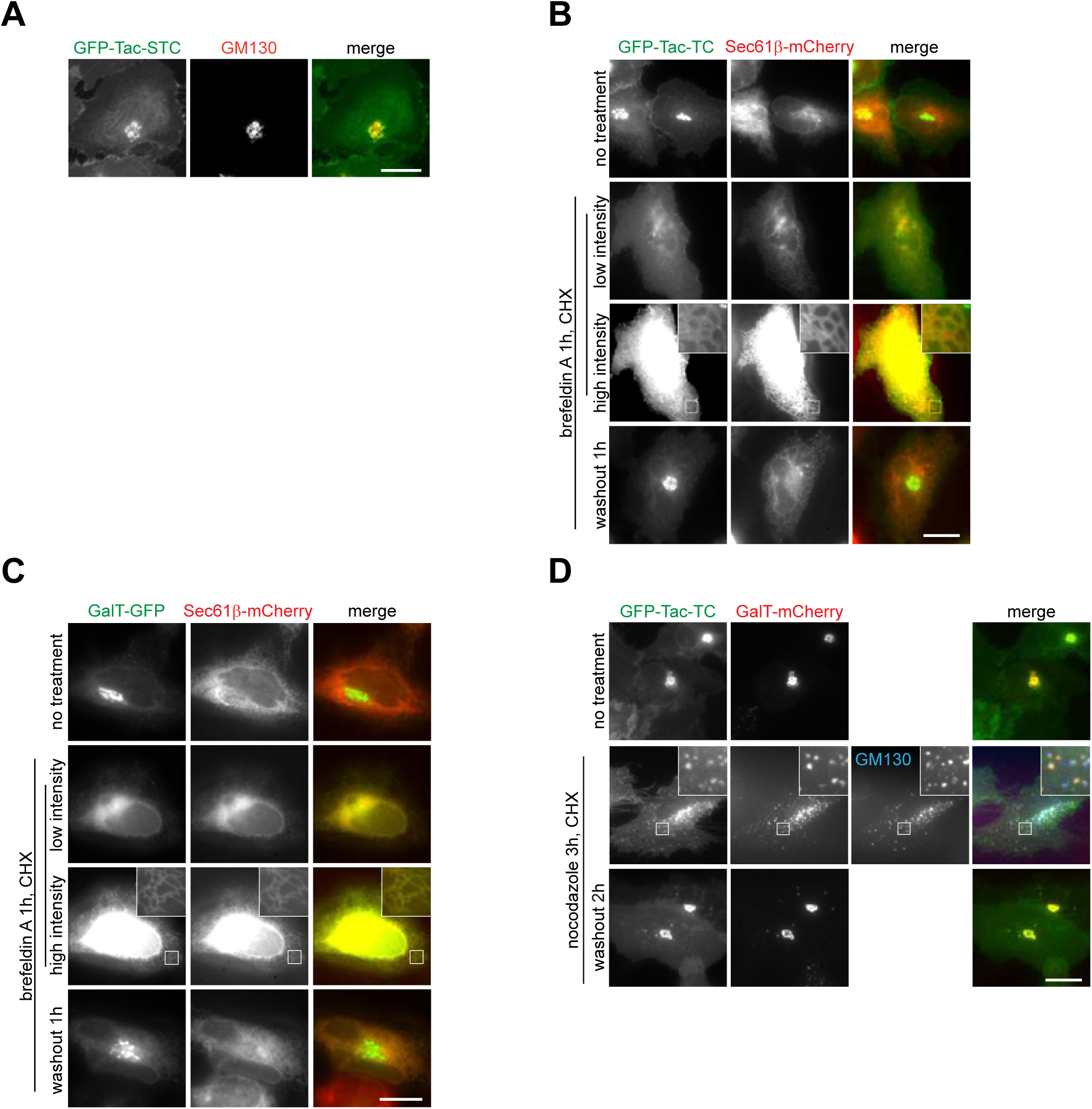
Tac-TC behaves like a Golgi resident. HeLa cells were used. (A) Example image showing the Golgi localization of GFP-Tac-STC. Cells transiently expressing GFP-Tac-STC were immuno-stained for endogenous GM130. (B,C) Tac-TC and GalT localize to the ER under brefeldin A treatment and to the recovered Golgi during the subsequent washout. Cells transiently co-expressing Sec61β-mCherry and GFP-Tac-TC (B) or GalT-GFP (C) were subjected to the indicated treatment and imaged live. The same images with different intensity scaling are shown in the second and third row to reveal the weak ER localization signal. (D) Tac-TC and GalT localize to the Golgi ministack under nocodazole treatment and to the recovered Golgi during the subsequent washout. Cells transiently co-expressing GFP-Tac-TC and GalT-mCherry were subjected to the indicated treatment and subsequently immuno-stained for endogenous GM130 (for images in the second row). In (B-D), boxed regions are enlarged at the upper right corner; scale bar, 20 µm.

**Supplementary Figure 2.**
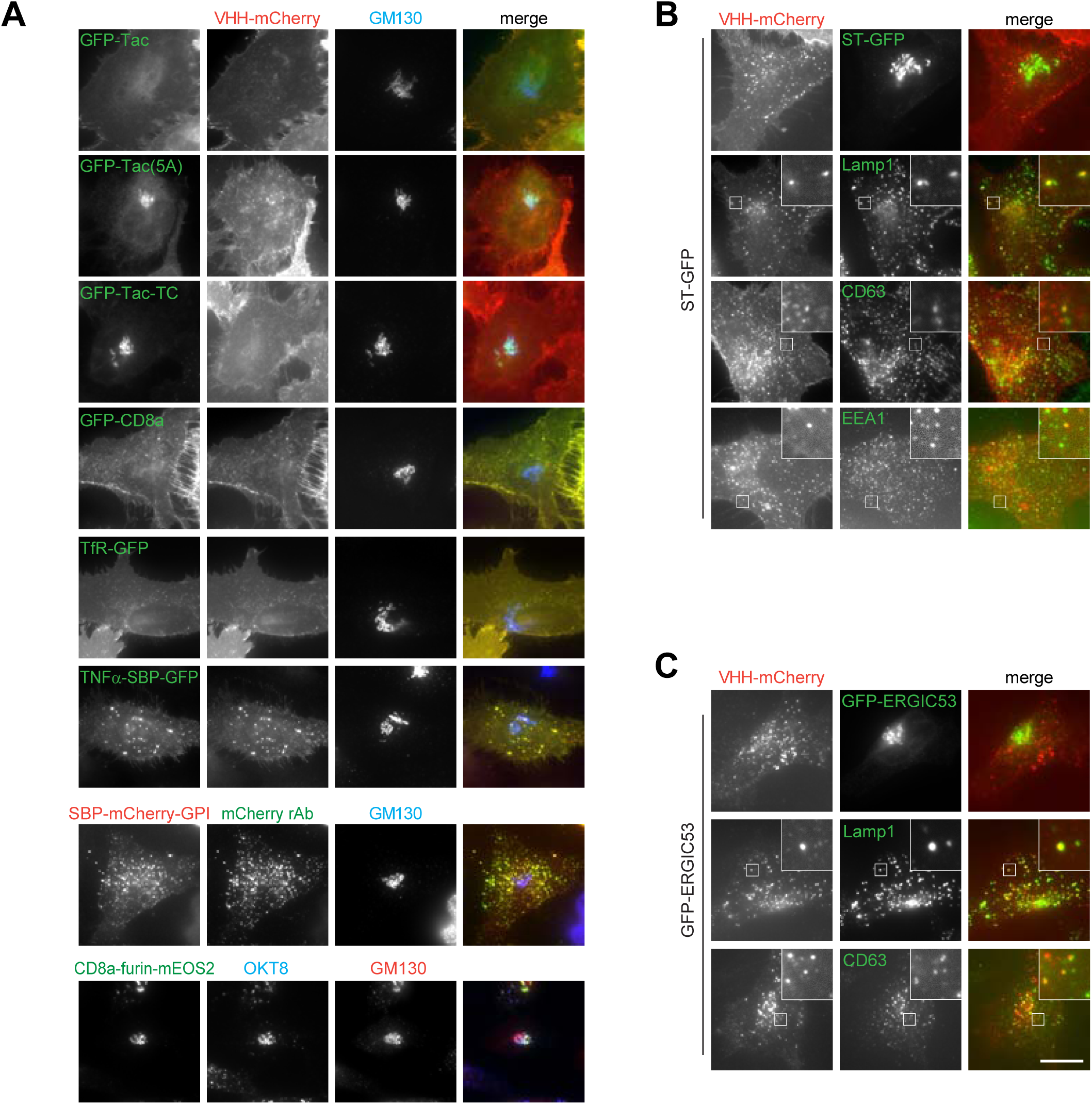
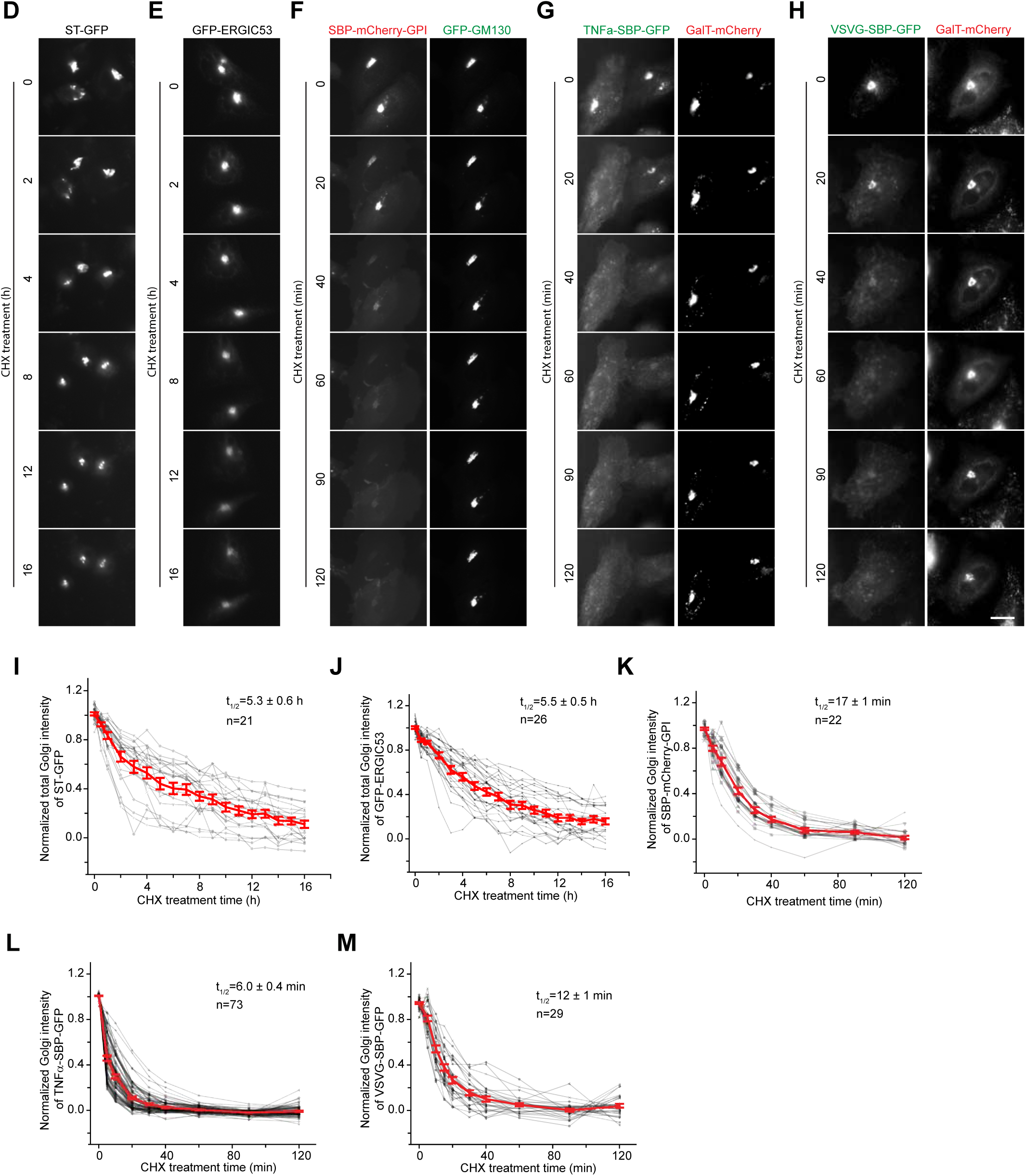
Investigating the endocytic trafficking and Golgi residence times of secretory cargos and Golgi residents. HeLa cells were used. (A-C) Reporters employed in this study do not target to the Golgi from the PM or endolysosome. In (A), cells transiently expressing indicated fluorescence protein-tagged reporter were incubated with VHH-mCherry, rabbit anti-mCherry polyclonal antibody or mouse anti-CD8a monoclonal antibody for 2 h and subsequently subjected to immuno-staining of endogenous GM130 and the internalized antibody. CD8a-furin-mEOS2 was a positive control for the Golgi targeting from the PM or endolysosome. In (B,C), surface-localized ST-GFP and GFP-ERGIC53 are targeted to the endolysosome instead of the Golgi. Cells transiently expressing ST-GFP (B) or GFP-ERGIC53 (C) were incubated with VHH-mCherry for 2 h and subsequently subjected to immuno-staining of endogenous Lamp1 (a lysosome marker), CD63 (a late endosome marker) or EEA1 (an early endosome marker). Boxed regions were enlarged at the upper right corner. (D-M) Acquiring the Golgi residence times of Golgi residents and secretory cargos. These panels are related to Figure 2D. In (D-H), cells expressing indicated fluorescence protein-tagged reporter(s) were imaged live in the presence of CHX. Although not shown, GalT-mCherry was also co-expressed in (D,E). In (I-M), the corresponding total Golgi intensity was plotted as described in Figure 2C. Scale bar, 20 µm; error bar, mean ± standard error; n, the number of quantified cells.

**Supplementary Figure 3.**
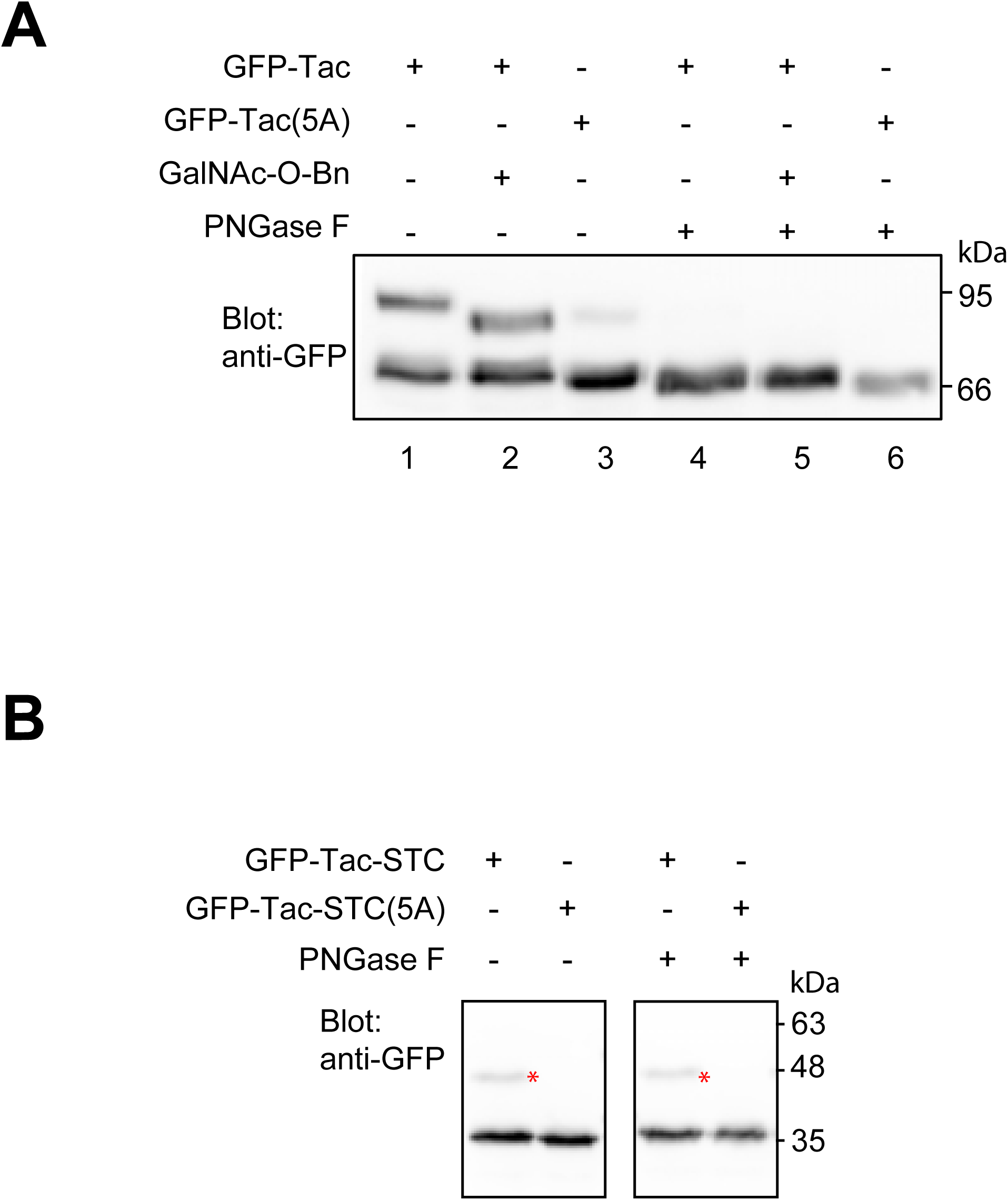
The gel migration profiles of Tac mutants. (A) GalNAc-O-Bn inhibits the O-glycosylation of Tac. HeLa cells transiently expressing GFP-Tac or GFP-Tac(5A) were treated or not with GalNAc-O-Bn for 20 h. Cell lysates were further treated or not with PNGase F before being immuno-blotted for GFP-tag. In comparison with lane 1, the disappearance of the upper band and the slightly increased migration of the lower band in lane 4 imply that the two bands of GFP-Tac in lane 1 are N-glycosylated. Likewise, lane 1 and 3 suggest that the two bands of GFP-Tac in lane 1 are O-glycosylated. The increased migration of the upper band in lane 2 compared with lane 1 indicates that GalNAc-O-Bn probably inhibits the O-glycosylation of Tac. (B) Only a small fraction of Tac-STC is O-glycosylated. The lysate of HeLa cells transiently expressing GFP-Tac-STC or GFP-Tac-STC(5A) was treated or not with PNGase F before immune-blotted for GFP-tag. * indicates the O-glycosylated band. The left and right panels are cropped from the same gel blot. Molecular weight markers (kDa) are indicated at the right.

**Supplementary Figure 4.**
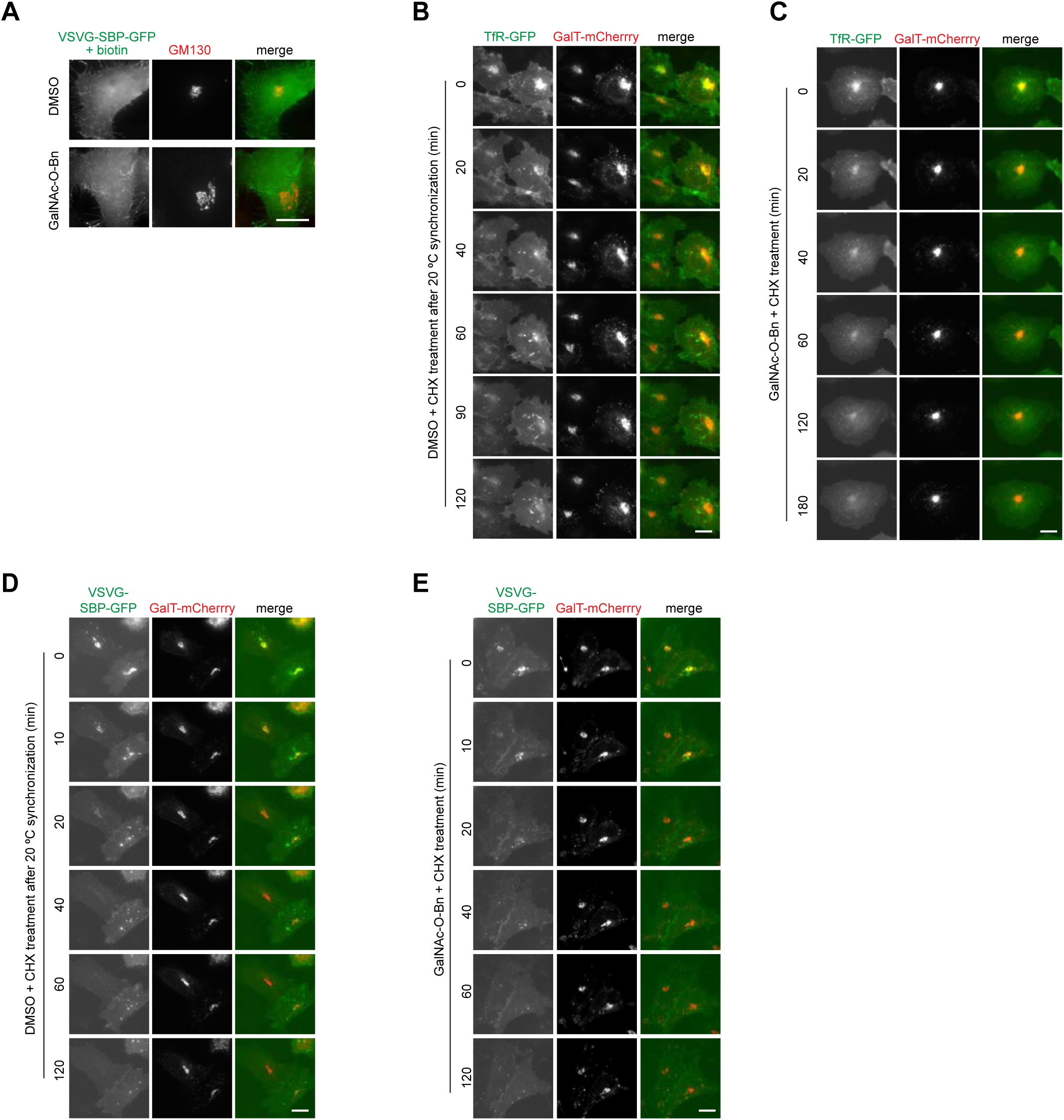
Inhibiting the O-glycosylation results in the substantial Golgi localization of TfR but not VSVG. HeLa cells were used. (A) After the treatment of DMSO or GalNAc-O-Bn, cells transiently expressing VSVG-SBP-GFP were further treated with biotin for 14 h before immuno-staining of endogenous GM130. (B,C) Cells transiently co-expressing TfR-GFP and GalT-mCherry were subjected to similar experimental procedure as described in Figure 4D. (D,E) As described in Figure 4I,J. Scale bar, 20 µm.

**Supplementary Figure 5.**
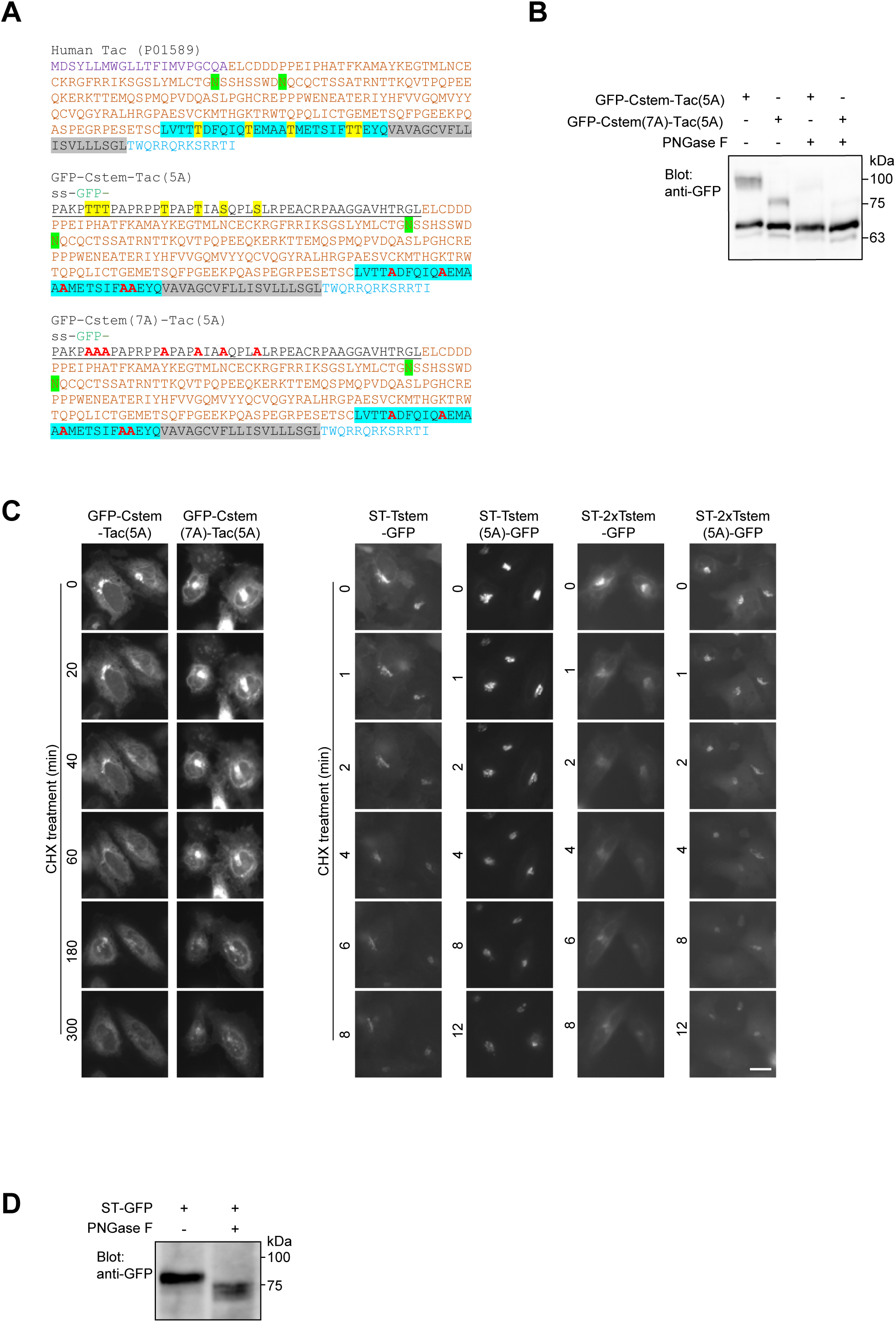
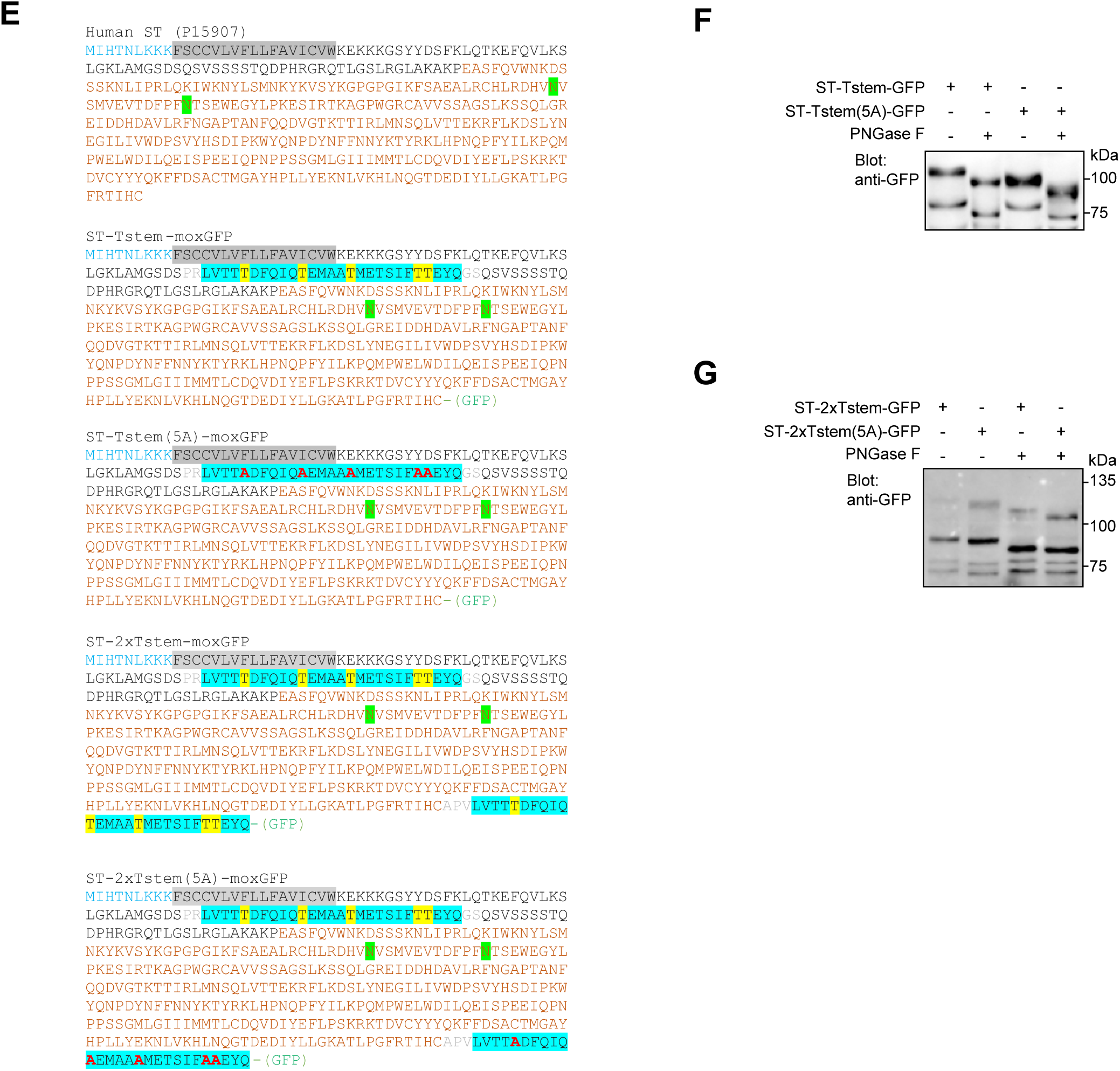
The sequences, gel migration profiles, and time-lapse images of Tac and ST chimeras. HeLa cells were used. (A) The sequences of Tac (Uniprot identifier: P01589) chimeras. Purple AAs, signal sequence (hereafter ss); blue AAs, cytosolic tail; grey shaded AAs, transmembrane domain; brown AAs, IBD; green shaded Ns, N-glycosylation sites; cyan shaded AAs, Tstem; underlined AAs, Cstem. In Tstem and Cstem, potentially O-glycosylated AAs (Thr or Ser) are shaded yellow and the corresponding Ala mutations are colored red. In chimera sequences, GFP is inserted between the ss and Cstem. (B) The gel migration profile of Tac chimeras demonstrates that Cstem likely undergoes the O-glycosylation. Lysates of cells transiently expressing indicated constructs were treated or not with PNGase F before immuno-blotting for GFP-tag. (C) The time-lapse images of Tac and ST chimeras. Cells transiently co-expressing GalT-mCherry and indicated construct were live-imaged in the presence of CHX. Scale bar, 20 µm. (D) The gel migration profile of ST demonstrates that it is likely N-glycosylated. The experimental procedure was similar to that of (B). (E) The sequences of ST (Uniprot identifier: P15907) chimeras. AAs are colored or shaded as described in (A) except that brown AAs indicate the catalytic domain. GFP is appended at the C-terminus of chimeras. (F,G) The gel migration profiles demonstrate that Tstem is likely glycosylated as designed in ST-Tstem-GFP and ST-2xTstem-GFP. Experiments were conducted as described in (B). In (B,D,F,G), molecular weight (kDa) is indicated at the right side.

**Supplementary Figure 6.**
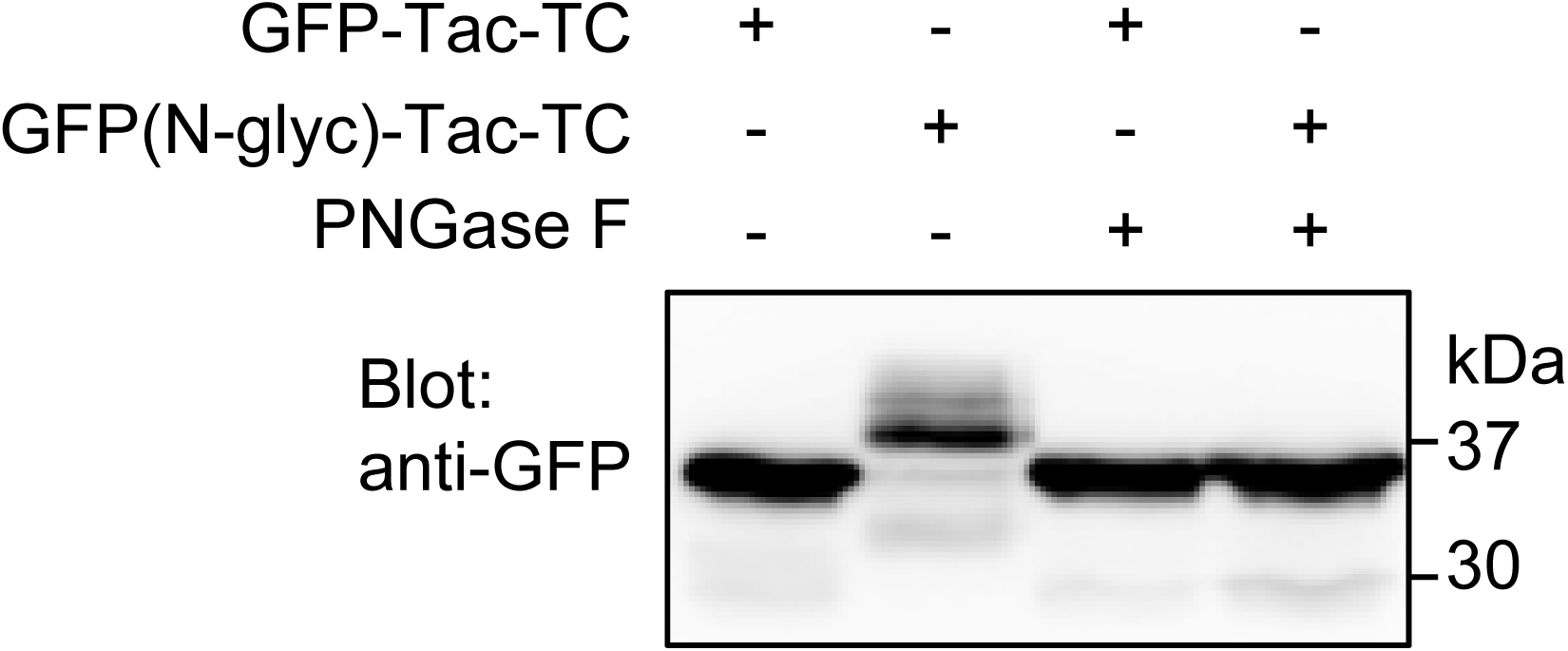
GFP(N-glyc)-Tac-TC is N-glycosylated. HeLa cells transiently expressing indicated construct were lysed and treated or not with PNGase F before gel separation and immuno-blotting for GFP-tag. Molecular weight (kDa) is indicated at the right side.

**Supplementary Figure 7.**
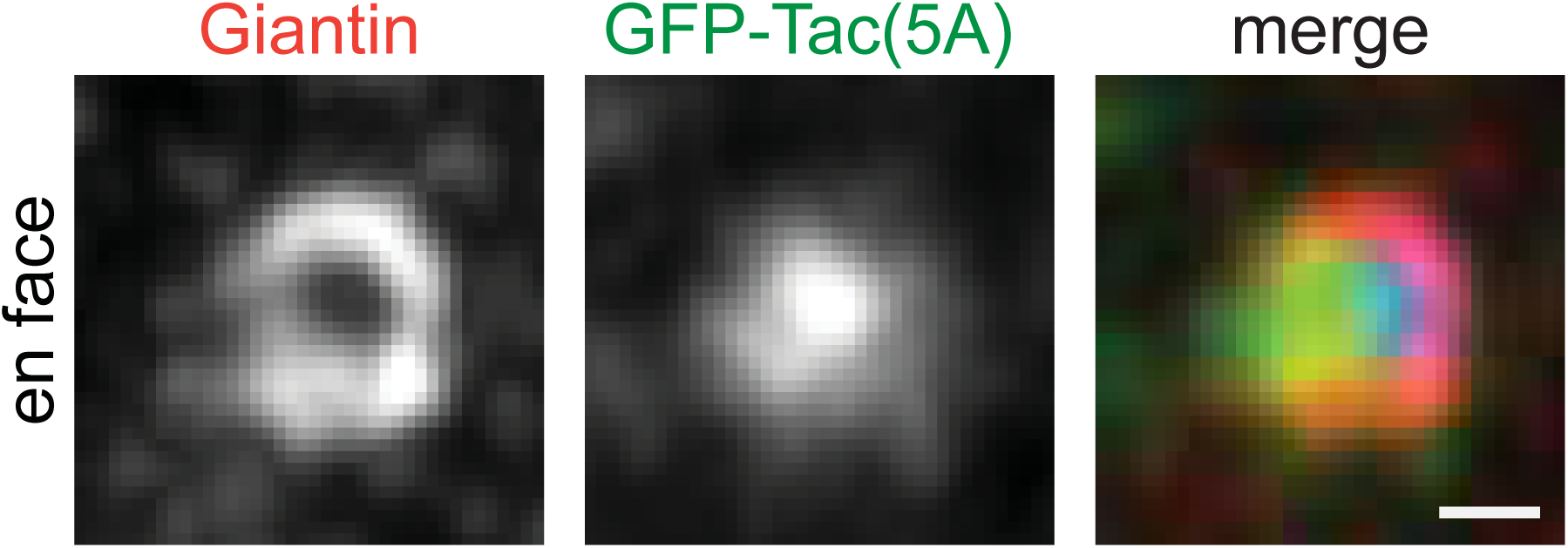
GFP-Tac(5A) localizes to the interior of the *trans*-Golgi cisternae. The experiment is similar to that of Figure 7D. Scale bar, 500 nm.

**Supplementary Table 1.**
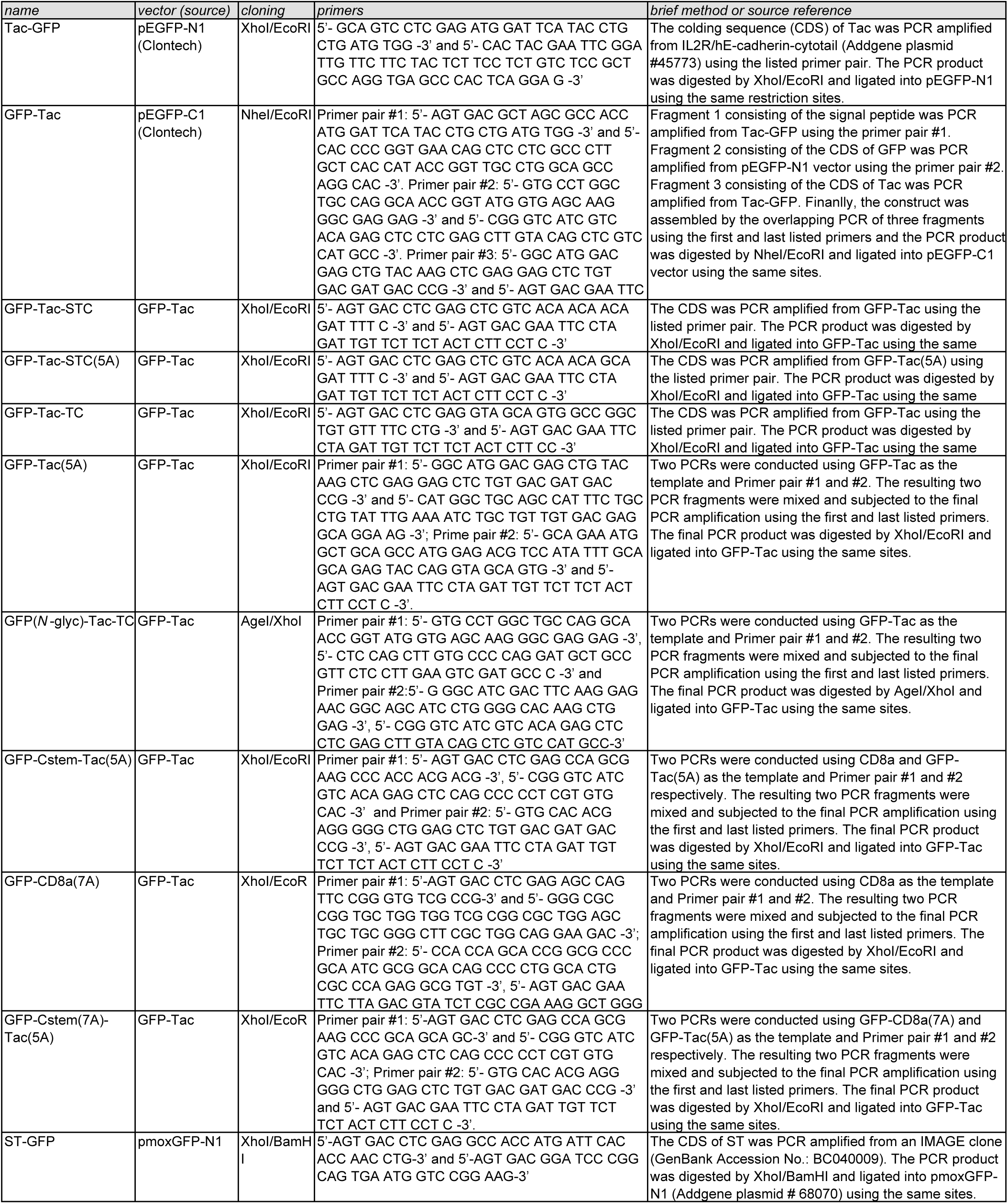
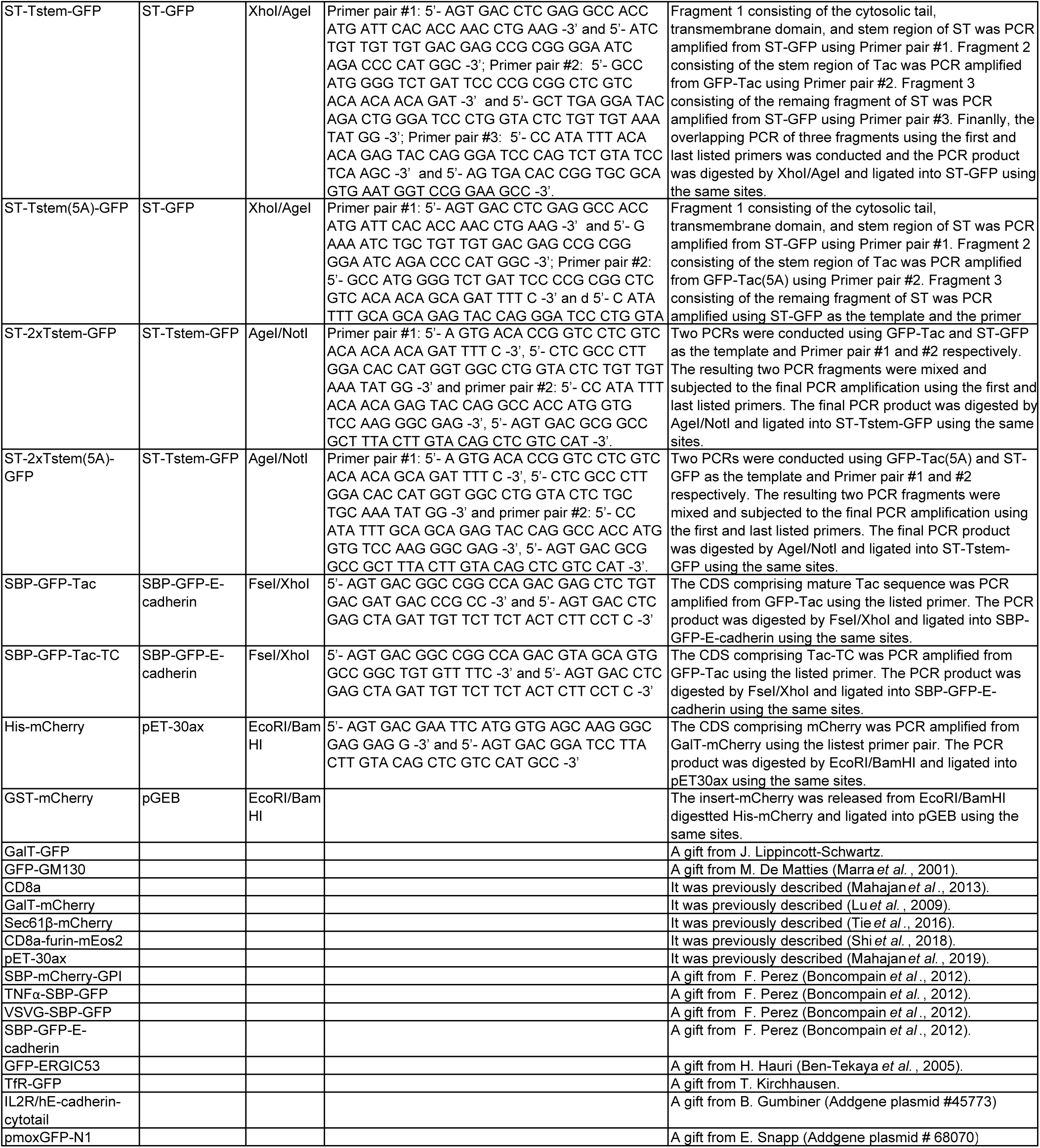
List of DNA plasmids.

## Notes

### Competing Interest Statement

The authors have declared no competing interest.

